# CENP-B binds hairpin motifs in chromosome arms influencing gene expression

**DOI:** 10.64898/2026.04.10.717743

**Authors:** Lillian Wu, Karen A Lane, Reyhan Muhammad, Alison Harrod, Catherine Naughton, Hefei Wang, Andrea Musacchio, Nick Gilbert, Claudio Alfieri, Jessica A Downs

## Abstract

CENP-B, a centromeric protein known for its role in binding the B box sequence of centromeric DNA, has long been recognized as important, though not essential, for kinetochore attachment and chromosome segregation. Here, we identify an unexpected, non-centromeric role for CENP-B. We demonstrate that CENP-B binds to specific non-centromeric sites along chromosome arms, predominantly at promoters, and depletion of CENP-B leads to dysregulated gene expression. Binding is enriched in G2 phase cells and, importantly, occurs independently of the canonical B box motif. Instead, CENP-B binding in chromosome arms is defined by regions of negatively supercoiled DNA containing repetitive sequences, such as multiple CCAAT boxes, that are prone to forming secondary structures. Consistently, we find that CENP-B binds to hairpin DNA *in vitro* via its DNA binding domain. The chromosome arm binding pattern is conserved across cell types and is particularly prominent in the promoters of transcriptionally active replication-dependent histone genes. These findings reveal a previously unrecognized centromere-independent binding activity of CENP-B.

## INTRODUCTION

CENP-B was discovered over 40 years ago^1^ and since then a great deal of work has contributed to understanding its role at centromeres ^2–4^. CENP-B interacts with CENP-A-containing nucleosomes and components of the constitutive centromere-associated network (CCAN) complex^4,5^. These complexes play a vital role mediating kinetochore attachments and promoting appropriate chromosome segregation.

CENP-B has an N-terminal DNA binding domain (DBD), a transposase-like domain, an acidic region, and a C-terminal dimerisation domain. The CENP-B DBD has been shown to bind a 17 bp consensus sequence found in centromeric repeats termed the B box^6,7^, and this binding contributes to fidelity of chromosome segregation^5^. Structural studies indicate that this interaction is mediated through major groove interactions with 9 bp within the B box^8^. Chromosomes undergo large-scale changes in organisation during the cell cycle to position centromeres at the chromatin surface, and recently, the acidic region of CENP-B was identified as a key player in this process^9^.

Interestingly, unlike CCAN complex subunits and CENP-A, CENP-B is not essential for centromere function^3^ and CENP-B knockout mice are viable^10–12^. While CENP-B homologues in fission yeast have been shown to silence transposable elements^13,14^ and promote replication through these sites^15^, non-centromeric functions have not been identified for CENP-B in other organisms, including humans.

## RESULTS

### CENP-B binds to gene regulatory sites in chromosome arms

To understand more about CENP-B function in human cells, we performed CENP-B mapping using CUT&RUN in asynchronous RPE1 cells. As expected, we found CENP-B localised to B-box motifs in centromeres (Fig. 1a and Supplementary Fig. 1a). Interestingly, we noticed that CENP-B also binds to a subset of genomic locations outside centromeres (Fig. 1b). We excluded centromere-associated enrichment and identified consensus non-centromeric CENP-B (hereafter referred to as nc-CENP-B) peaks found in all three biological replicates. By overlapping these locations with ATAC-seq analysis performed in the same cells, we find that these are located at sites of open chromatin (Fig. 1 and Supplementary Fig. 1b) and are predominantly found at gene promoters (Fig. 1d,e and Supplementary Fig. 1c). Pathway analysis of the gene promoters with nc-CENP-B show enrichment of cell cycle gene sets (Supplementary Fig. 1d). These data raise the possibility that CENP-B might play a role in the expression of a subset of genes.

**Figure 1:**
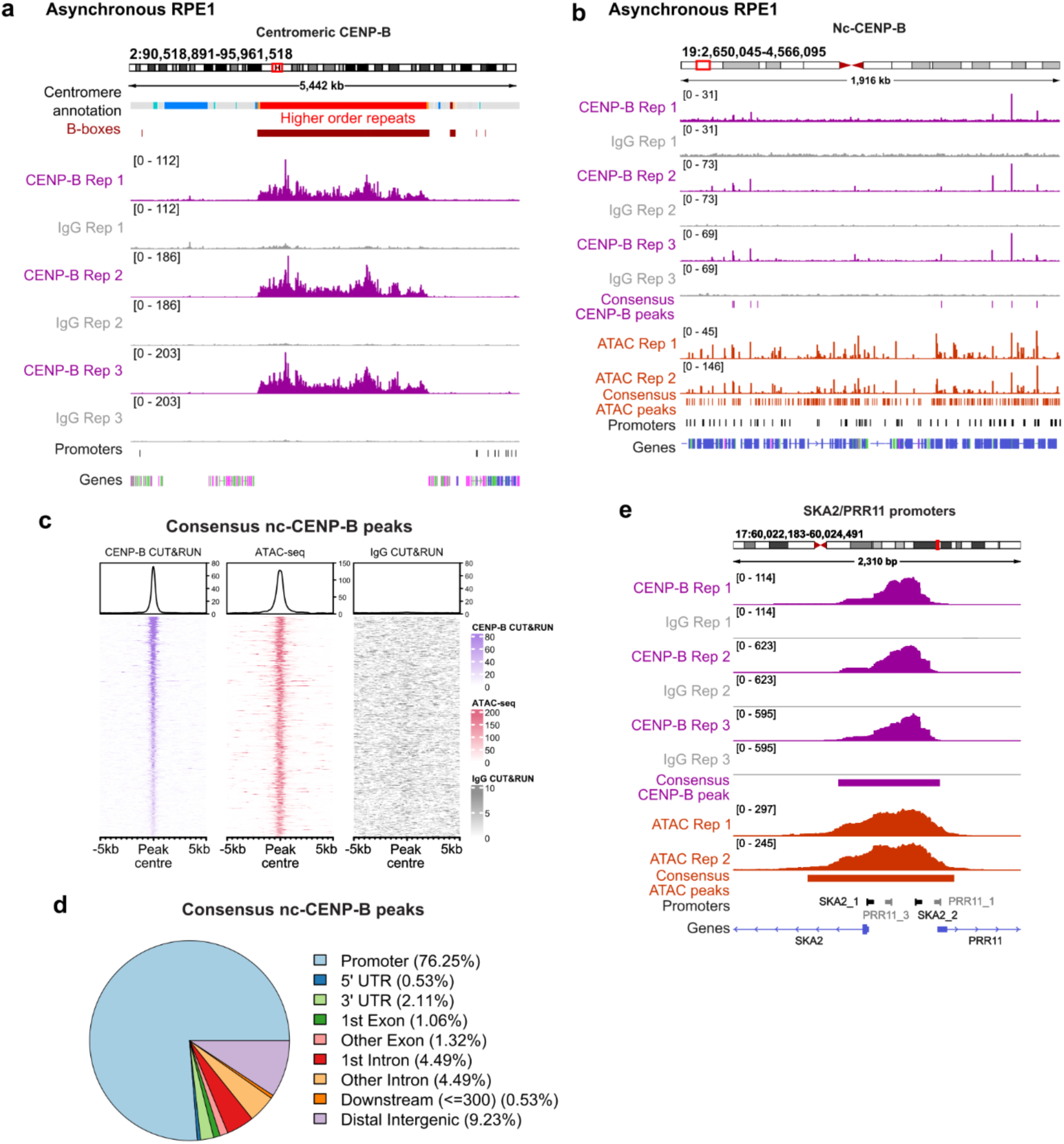
Non-centromeric CENP-B (nc-CENP-B) binds to gene regulatory sites in chromosome arms. **(a)** IGV screenshot showing CENP-B binding across the centromere of chromosome 2 in RPE1 cells, showing coverage of reads from CENP-B (purple) and IgG (grey) CUT&RUN in 3 biological replicates. Locations of active higher order repeats (HORs) are annotated in red, and canonical B-box motifs in brown. **(b)** IGV screenshot showing CENP-B binding sites along the chromosome arm of chromosome 19, showing coverage of reads from 3 biological replicates of CENP-B (purple) and IgG (grey) CUT&RUN sequencing in RPE1 cells. ATAC-seq read coverage in RPE1 cells from 2 biological replicates are also shown (orange). Gene promoters and transcripts are indicated below. **(c)** Heatmap showing CENP-B CUT&RUN (purple), ATAC-seq (red), and IgG CUT&RUN (grey) signal intensity at consensus nc-CENP-B peaks in 3 of 3 biological replicates (379 peaks), showing regions +/-5kb of each peak centre. An average CUT&RUN or ATAC-seq signal plot is displayed above. **(d)** Pie chart showing genomic features occupied by the consensus RPE1 nc-CENP-B peaks, indicating the percentages of peaks within each type of feature. Promoters = 1 kb upstream and 0.2 kb downstream of the TSS. **(e)** IGV screenshot showing CENP-B binding at the SKA2 and PRR11 gene promoters, showing coverage of reads from CENP-B (purple) and IgG (grey) CUT&RUN sequencing, and ATAC-seq read coverage in orange. The locations of the SKA2 and PRR11 promoters and transcripts are shown below.

We set out to test this using RNA-seq following CENP-B depletion. However, we found that depleting CENP-B in asynchronous cells altered the cell cycle profile (Supplementary Fig. 2a). Since many of the nc-CENP-B-bound promoters are cell cycle regulated genes, this could confound interpretation. To avoid this issue, we depleted CENP-B in cells arrested in G1 phase using palbociclib or in G2 phase using RO-3306 (Fig. 2a and Supplementary Fig. 2a,b). Using these conditions, we performed RNA-seq and found that depletion of CENP-B led to altered gene expression – both up-and downregulation – in both phases of the cell cycle (Fig. 2b).

**Figure 2:**
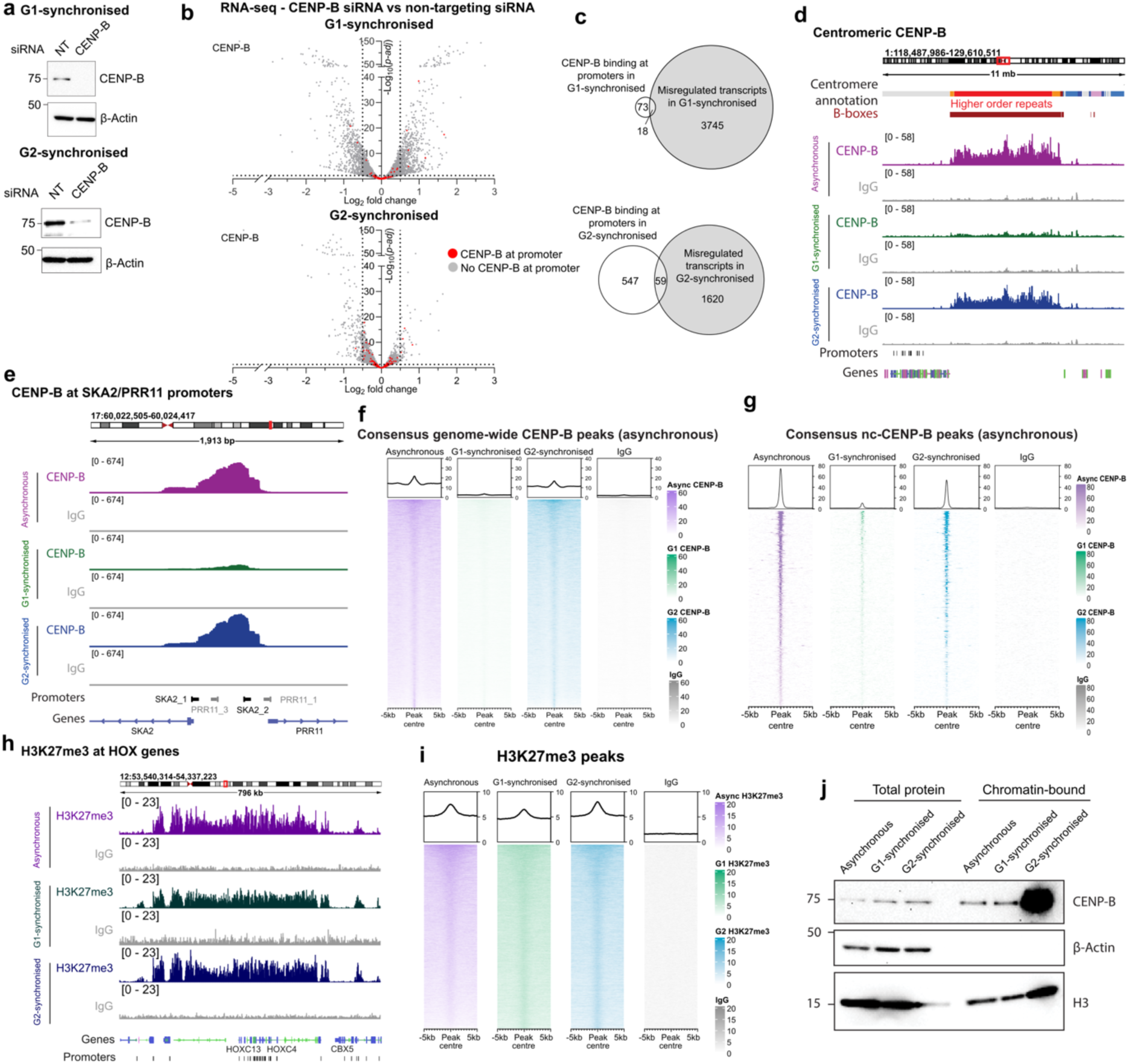
CENP-B influences gene expression. **(a)** Representative Western Blots showing siRNA-mediated CENP-B depletion (and non-targeting siRNA control) in G1 and G2-synchronised RPE1 cells. **(b)** Volcano plots showing expression changes of all transcripts in CENP-B siRNA-treated compared to non-targeting siRNA-treated cells, for G1 (n = 3) and G2-synchronised (n = 2) cells. The transcript log2 fold change (x-axis) and-log10(p-adj) (y-axis) are shown, genes with CENP-B binding at their promoter are in red. **(c)** Venn diagrams showing the number of overlaps between expressed genes with expression changes (base mean > 10 and p-adj < 0.05) after CENP-B siRNA treatment (vs non-targeting control) in G1/G2-asynchronised cells and genes with CENP-B binding at promoters in G1/G2-synchronised cells. The size of each circle and of the overlapping region is proportional to the number of genes. **(d-e)** IGV screenshots showing coverage of CENP-B and IgG CUT&RUN reads in asynchronous (purple), G1-synchronised (green), G2-synchronised (blue) RPE1 cells across the centromere of chromosome 1 (d) and in the arm of chromosome 9 (e). **(f-g)** Heatmaps of CENP-B CUT&RUN signal intensity in asynchronous (purple), G1-synchronised (green), and G2-synchronised (blue) cells at consensus genome-wide CENP-B peaks (g) and consensus nc-CENP-B peaks (h) in asynchronous cells. An average CUT&RUN signal plot is displayed above. **(h)** IGV screenshot showing coverage of H3K27me3 and IgG CUT&RUN reads in asynchronous (purple), G1-synchonised (green), G2-synchronised (blue) cells across the HOXC genes cluster. **(i)** Heatmaps of H3K27me3 CUT&RUN signal intensity in asynchronous (purple), G1-synchronised (green), and G2-synchronised (blue) cells at subsampled (n = 10,000) consensus H3K27me3 peaks in G2-syncrhonised cells. **(j)** Western Blot showing levels of total and chromatin-bound CENP-B in asynchronous, G1-synchronised, and G2-synchronised RPE1 cells, with B-actin and histone H3 loading controls.

We also mapped CENP-B in G1 and G2 phase cells to directly compare CENP-B promoter binding with CENP-B-dependent gene regulation. Surprisingly, very few of the genes whose expression changes when CENP-B is depleted also had CENP-B associated with their promoter, and conversely, most genes with CENP-B occupancy at their promoter were not significantly misregulated when CENP-B is depleted (Fig. 2b,c). Interestingly, however, pathway analysis of the misregulated genes shows an enrichment of cell cycle regulated genes in G2 phase cells (Supplementary Fig. 2c), which is similar to the pathway analysis of genes that have CENP-B bound at their promoter, raising the possibility of a functional relationship.

Analysis of these data revealed that CENP-B enrichment – both at centromeres and in chromosome arms – is much stronger in G2 phase than G1 (Fig. 2d-g and Supplementary Fig. 2d-g). In contrast, H3K27me3 enrichment exhibited minimal variation between G1 and G2 phase samples (Fig. 2h,i). Notably, CENP-B signals remained elevated in G2 phase samples even after normalisation with a spike-in control (Supplementary Fig. 2h,i). To explore this further, we performed chromatin fractionation and found that chromatin-associated CENP-B levels are substantially greater in G2 phase cells (Fig. 2j). These data are consistent with a previous report showing that the exchange rate of CENP-B on chromatin is much greater in G1 compared with G2^16^.

### Sequence and structural determinants of nc-CENP-B binding

B box motifs are not exclusively centromeric and exist in chromosome arms^17^, raising the possibility that nc-CENP-B is directed to chromosome arms through binding to this motif. Surprisingly, however, motif analysis of nc-CENP-B peaks shows that these sites are not enriched for B boxes, but instead are most highly enriched for CCAAT boxes (Fig. 3a). Looking at these sites in more detail, we find that nc-CENP-B sites tend to have multiple CCAAT boxes, more than expected by chance, and often in both orientations forming inverted repeats (Fig. 3b and Supplementary Fig. 3a,b).

**Figure 3:**
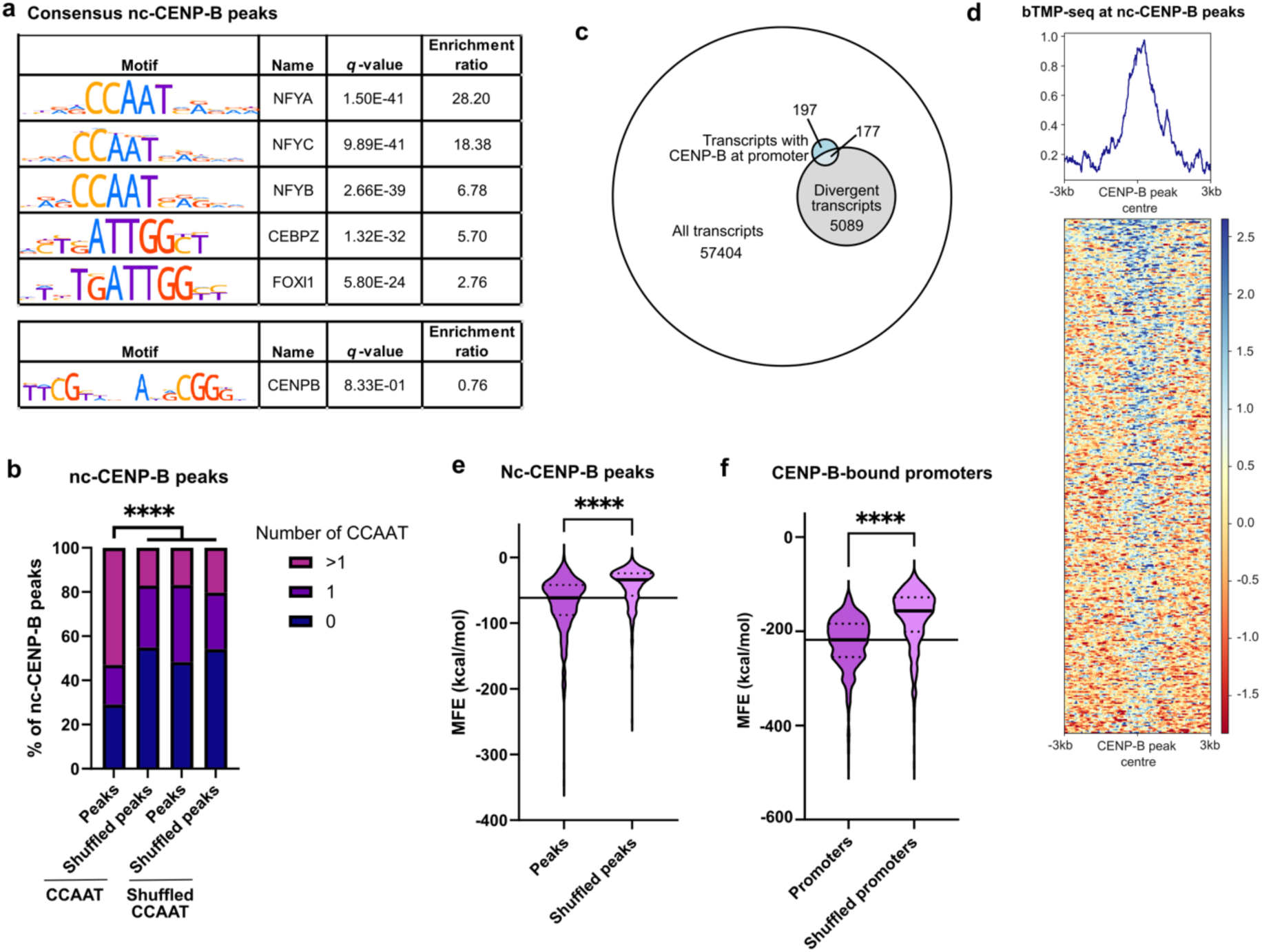
nc-CENP-B binds CCAAT boxes and secondary structure-forming sequences. **(a)** Motif enrichment analysis results for consensus nc-CENP-B peaks in asynchronous cells, showing enrichment of the CCAAT box-containing motifs (top) and the B-box motif (bottom). Enrichment ratio indicates the relative enrichment of each motif in the DNA sequences of nc-CENP-B peaks compared to shuffled input sequences. **(b)** Bar chart showing the percentages of CENP-B peaks containing the specified number of CCAAT boxes and shuffled CCAAT box coordinates for nc-CENP-B peaks and shuffled nc-CENP-B peaks. Data were analysed by Chi-square test (**** p < 0.0001), comparing count distribution in bar 1 to the average count distribution across bars 2-4 (negative controls). **(c)** Venn diagram showing the number of bidirectional promoters (Divergent transcripts) out of all annotated GENCODE promoters, and the number of promoters occupied by CENP-B. Numbers denote the number of transcripts in each category in the Venn diagram. **(d)** Heatmap showing bTMP-seq signal (negative supercoiling) intensity at consensus non-centromere CENP-B peaks, showing regions +/-3 kb of each CENP-B peak. An average bTMP-seq signal plot is shown. **(e)** Violin plots of minimum free energy (MFE) predictions for RPE1 consensus nc-CENP-B peaks, and shuffled nc-CENP-B peaks. **(f)** Violin plots of MFE predictions for promoters bound by CENP-B, and shuffled set of promoters. For e-f, data were analysed by two-tailed Welch’s t-test (**** p < 0.0001), the median is marked by a solid line, and quartiles are marked by dashed lines.

We noticed a disproportionate representation of bidirectional promoters with divergently transcribed genes in the nc-CENP-B peaks. While divergent transcripts comprise less than ten percent of all transcripts, almost half of the transcripts with CENP-B at the promoter are found at these sites (Fig. 3c). Gene expression at bidirectional promoters can lead to increased negative supercoiling, which in turn may stabilise DNA secondary structure formation when inverted repeats are present^18,19^. We therefore asked if the enrichment of CENP-B at these promoters may reflect a preference for supercoiled, and consequently structured, DNA. To do this, we interrogated the supercoiling landscape using biotynlated-trimethylpsoralen sequencing (bTMP-seq). We found a correlation between nc-CENP-B peaks and negative supercoiling (Fig. 3d).

Given the propensity of CENP-B occupied sites to have both CCAAT box repeats and negative supercoiling, these observations raise the possibility that nc-CENP-B might be enriched at secondary structure-forming sites, which, notably, is also a feature of centromeric DNA^20,21^ (Supplementary Fig. 3c). In support of this, we find that the nc-CENP-B peaks have a lower in silico secondary structure minimum free energy (MFE) score than expected by chance (Fig. 3e), which indicates an increased likelihood of secondary structure formation. This is even more striking when restricting analysis to promoter-bound nc-CENP-B (Fig. 3f).

### CENP-B directly binds to hairpin-containing DNA *in vitro* via its DNA binding domain

To test whether CENP-B is directly capable of binding to sequences found in the nc-CENP-B enriched peaks, we performed *in vitro* pull-down assays with recombinant CENP-B and a series of oligos (Fig. 4a,b). As expected, CENP-B bound to double-stranded DNA (dsDNA) oligos containing a B box motif in preference to dsDNA with a shuffled B box sequence (Fig. 4c). Using dsDNA oligos containing either 2 or 4 CCAAT boxes, we found no detectable binding above background levels (Fig. 4c).

**Figure 4:**
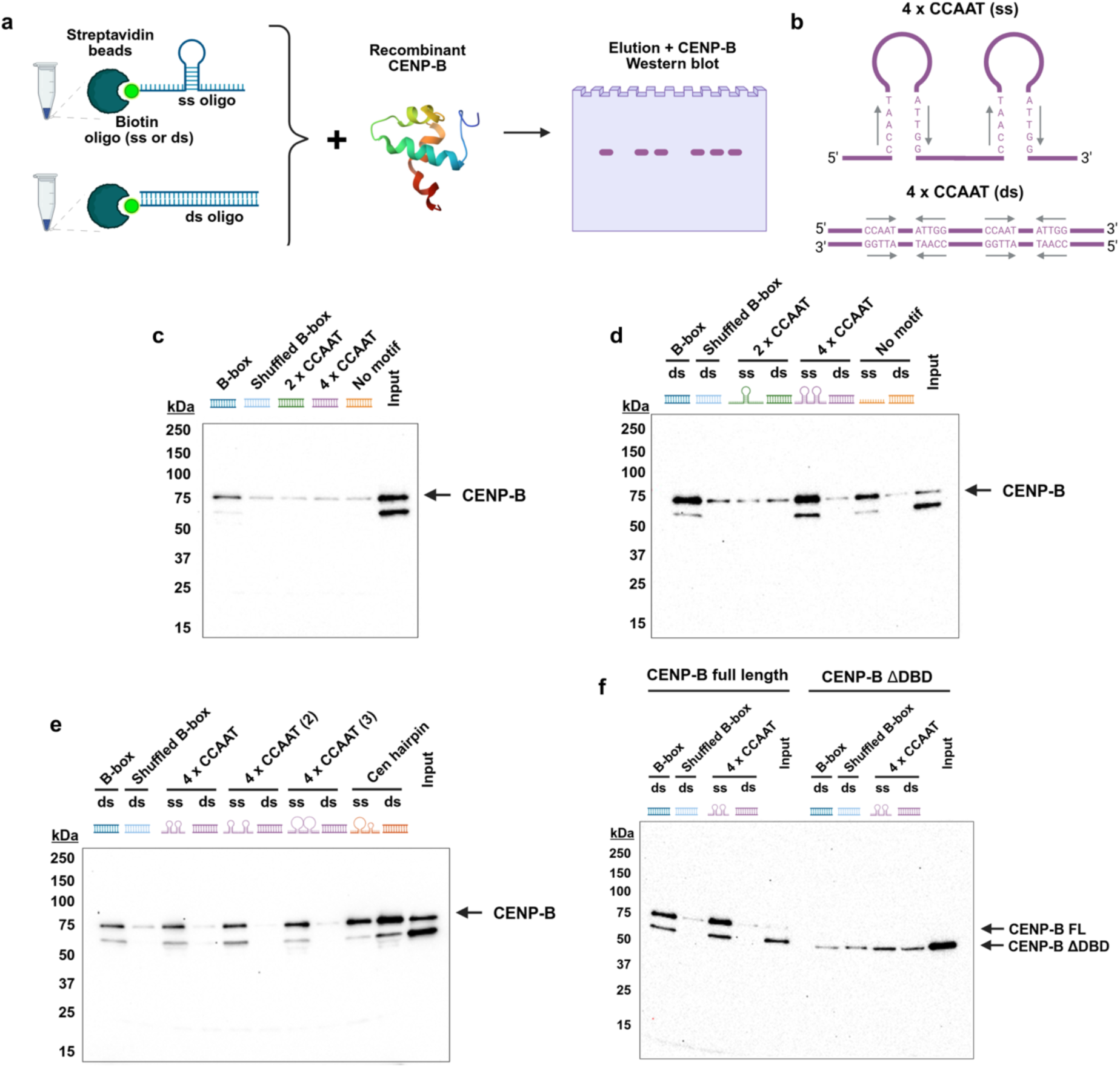
CENP-B binds CCAAT-containing hairpin-forming sequences *in vitro*. **(a)** Schematic of CENP-B pull-down experiments: biotinylated DNA oligos are immobolised on Streptavidin beads, then recombinant CENP-B is added and incubated. DNA-bound CENP-B is eluted and visualised by Western blotting **(b)** Schematics showing the structure and CCAAT boxes orientations for the 4 x CCAAT ssDNA and dsDNA oligos. **(c)** Results of *in vitro* pull-down assays showing recombinant full-length CENP-B binding to different dsDNA oligos annotated above. **(d-e)** Results of *in vitro* pull-down assays showing recombinant full-length CENP-B binding to different ssDNA and dsDNA oligos annotated above. **(f)** *In vitro* pull-down assays showing recombinant CENP-B and CENP-B ΔDBD binding to different DNA oligos annotated above. Figures were generated using BioRender (www.biorender.com)

We therefore tested whether CENP-B is able to interact with the single-stranded DNA (ssDNA) CCAAT box-containing oligos when folded into hairpins. Strikingly, we found that CENP-B binds to the 4 x CCAAT box-containing ssDNA oligo, which forms two hairpins, to a similar level as the B box-containing dsDNA oligo (Fig. 4d). In contrast, CENP-B interacted less well with a ssDNA oligo forming a single hairpin or a ssDNA oligo without hairpin-forming sequence (Fig. 4d). Next, we varied the position of the inverted repeats to increase either the distance between the two hairpins or the length of the loop, but this had no obvious impact on CENP-B binding (Fig. 4e), suggesting that these features are not critical for the interaction. We also found that a hairpin-forming sequence from the centromere interacts with CENP-B (Fig. 4e), suggesting that the CCAAT box sequence is not the determining factor for hairpin binding.

To determine whether this interaction is mediated by the DNA binding domain (DBD) of CENP-B, we created a construct in which the DBD was removed and tested it in the pull-down assays. This construct was no longer able to interact with either the B box-containing dsDNA oligos or the CCAAT box-containing hairpin ssDNA oligos (Fig. 4f), suggesting that the interaction with both B box motifs and hairpin DNA is mediated by the DBD.

To test this further, we created a CENP-B expression construct lacking the DBD and expressed this or a full-length equivalent and performed CUT&RUN mapping in these cell lines (Supplementary Fig. 4a,b). As expected, truncation of the DBD leads to reduced enrichment at centromeres (Supplementary Fig. 4c), but we also find loss of CENP-B in chromosome arms (Supplementary Fig. 4d-f). Together with the *in vitro* binding assays, these data suggest that interaction between CENP-B and non-centromeric sites is mediated by the DBD directly binding to secondary structure forming sequences.

### Conservation of nc-CENP-B chromatin localisation across cell types

To determine whether this activity of CENP-B extends to other cell types, we made use of a CENP-B mapping dataset generated in the K562 chronic myelogenous leukaemia cell line^22^. We also mapped CENP-B in the embryonic kidney-derived HEK293 cell line. CENP-B is enriched at centromeres in all three cell lines as expected (Supplementary Fig. 5a). Interestingly, we also find a population of CENP-B bound to specific sites in the chromosome arms (Fig. 5a-c and Supplementary Fig 5b-d). Similar to nc-CENP-B in RPE1 cells, the majority of the nc-CENP-B peaks in K562 and HEK293 cells are associated with promoters (Fig. 5d).

**Figure 5:**
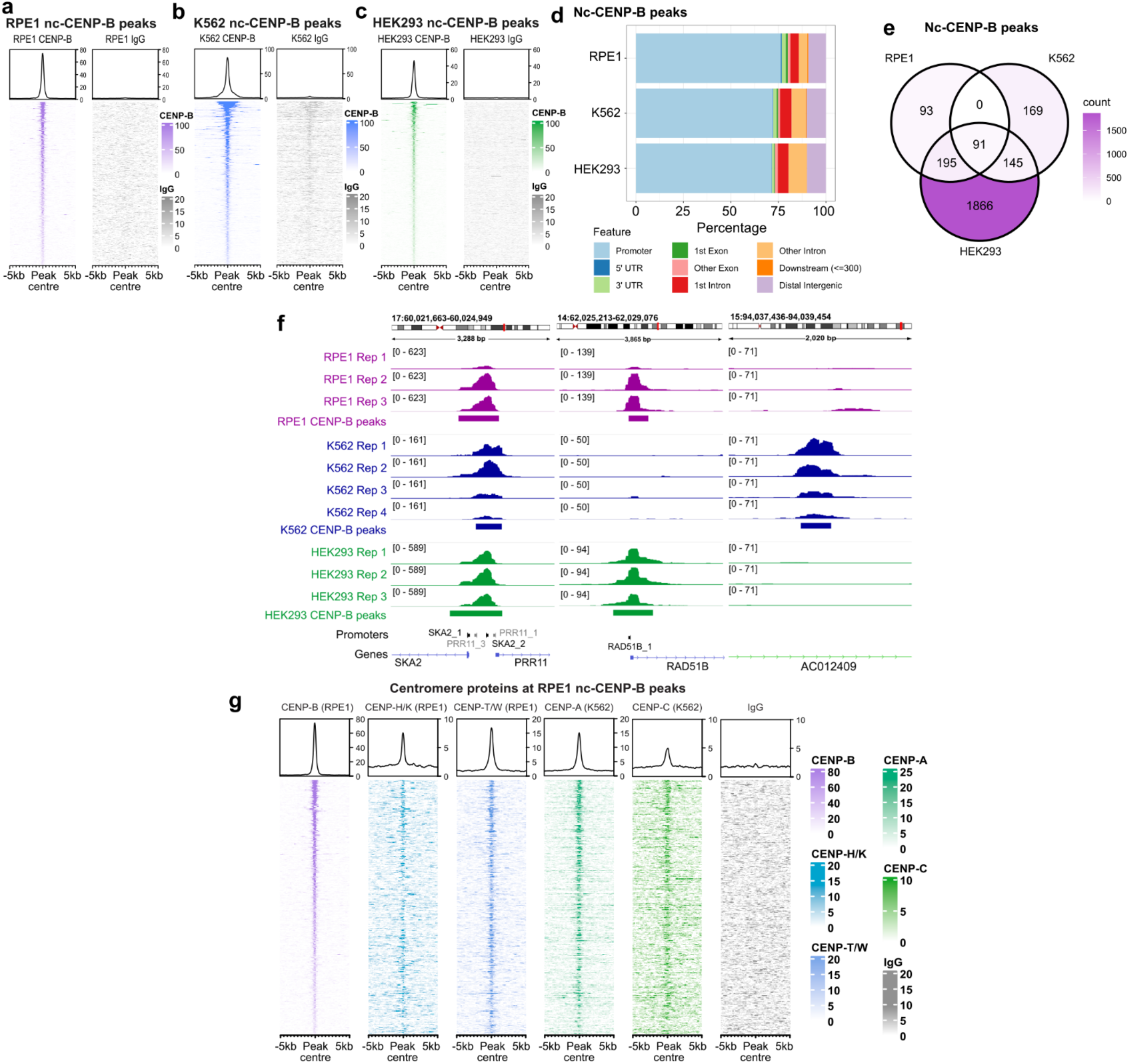
nc-CENP-B localisation is conserved across cell types. **(a)** Heatmaps of CUT&RUN signal intensity for CENP-B and IgG at consensus nc-CENP-B peaks in RPE1. **(b)** Heatmaps of CUT&RUN signal intensity for CENP-B and IgG at consensus nc-CENP-B peaks in K562. **(c)** Heatmaps of CUT&RUN signal intensity for CENP-B and IgG at consensus nc-CENP-B peaks in HEK293. **(d)** Bar chart showing genomic features occupied by the consensus nc-CENP-B peaks in RPE1, K562, and HEK293 cells, indicating the percentages of peaks within each type of feature. Promoters = 1 kb upstream and 0.2 kb downstream of the TSS. **(e)** Venn diagram showing the number of overlaps between consensus nc-CENP-B peaks in RPE1 (3 out of 3 replicates), K562 (4 out of 4 replicates), and HEK293 (3 out of 3 replicates). The colour corresponds to the total number of peaks in each region of the Venn diagram (count). **(f)** IGV screenshots showing coverage of CENP-B CUT&RUN reads in RPE1 (purple), K562 (blue), HEK293 (green) at the SKA2/PRR11 promoters, RAD51B promoter, and AC012409 genic region. **(g)** Heatmaps of CUT&RUN signal intensity for CENP-B in RPE1 (purple), CENP-H/K in RPE1 (light blue), CENP-T/W in RPE1 (dark blue), CENP-A in K562 (dark green), CENP-C in K562 (light green), IgG in RPE1 (grey) at consensus nc-CENP-B peaks in RPE1. An average CUT&RUN signal plot is displayed above each heatmap.

While there are peaks unique to each cell line, many nc-CENP-B sites are found in at least two of the three cell lines (Fig. 5e,f and Supplementary Fig. 5b-d and 6a-c), suggesting there is conservation in binding patterns.

In support of this, an analysis of the promoters bound in each cell line shows very similar enrichment of cell cycle related pathways (Supplementary Fig 6d,e). More compellingly, motif analysis of nc-CENP-B from K562 and HEK293 cells also shows that the CCAAT box is the most enriched motif (Supplementary Fig. 7a). Furthermore, nc-CENP-B sites in K562 and HEK293 cells tend to have multiple CCAAT boxes per binding site (Supplementary Fig. 7b) and have more stable secondary structure than expected by chance (Supplementary Fig. 7c,d). Together, these data suggest that the features of nc-CENP-B binding are conserved across cell types.

We next tested whether other centromere-associated proteins are present at nc-CENP-B bound sites in chromosome arms. To do this, we performed mapping in RPE1 cells using antibodies specific to CENP-H/K or CENP-T/W dimers, and we found enrichment at centromeres as expected (Supplementary Fig. 8a). Interestingly, we find there is detectable enrichment of these proteins at nc-CENP-B sites (Fig. 5g and Supplementary Fig. 8b), albeit at lower levels than CENP-B. We also mined CENP-A and CENP-C mapping data from K562 cells^22^ and find these are also present at nc-CENP-B sites (Fig. 5g and Supplementary Fig. 8a,b), suggesting these proteins might assemble at defined locations in chromosome arms.

### nc-CENP-B associates with actively transcribing replication-dependent histone genes

To further explore the association of CENP-B with chromosome arms, we plotted a) locations of genes with nc-CENP-B bound at their promoters and b) genes whose expression is affected by CENP-B depletion (Fig. 6a) and used this to determine regions where there is a significantly higher density of either of these than expected. Clusters of nc-CENP-B are apparent, as well as clusters of CENP-B-dependent genes. Several sites with high densities of both nc-CENP-B promoter binding and CENP-B-dependent gene expression are apparent, suggesting that perhaps clusters of nc-CENP-B bound sites influence gene expression in these broad regions.

**Figure 6:**
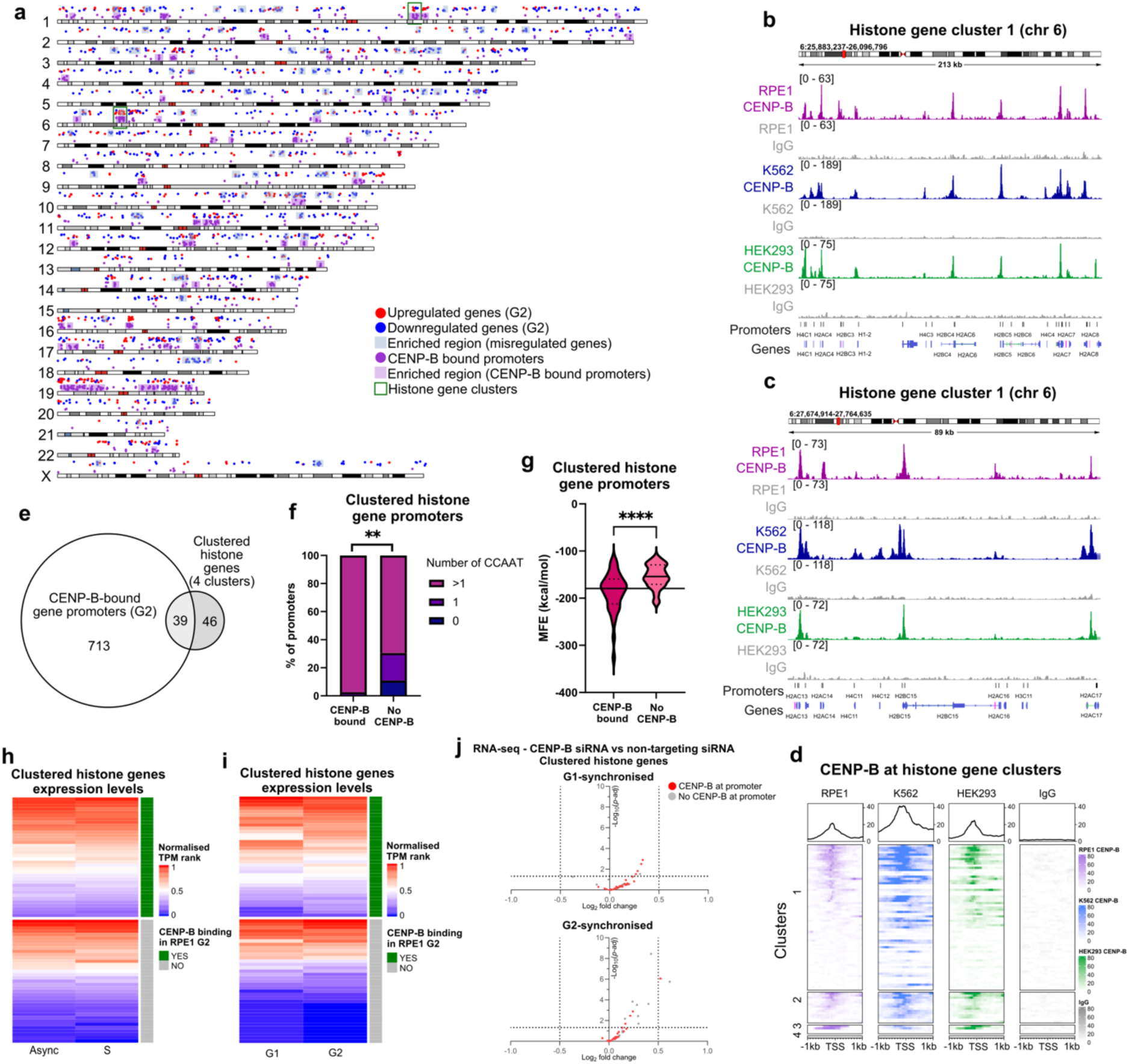
CENP-B binds histone genes and influences their expression. **(a)** Genome plots showing the locations of up-or down-regulated (p-adj < 0.05) genes in G2-synchronised cells treated with CENP-B-targeting versus non-targeting siRNAs (red or blue dots, respectively), and genes with CENP-B at their promoters in G2 (purple dots). The highlighted regions indicate clusters where there are a significantly enriched density (> 2 SD) of genes present in the set compared to all genes in the genome. Green boxes indicate locations of histone gene clusters 1 (Chr 6) and 2 (Chr 1). **(b-c)** IGV screenshots showing coverage of CENP-B and IgG CUT&RUN reads in RPE1 (purple), K562 (blue), and HEK293 (green) cells at two regions of histone gene cluster 1 in chromosome 6. **(d)** Heatmaps of CENP-B CUT&RUN signal intensity in RPE1 (purple), K562 (blue), and HEK293 (green) cells +/-1 kb from the TSS of the histone genes within the 4 histone gene clusters. An average CUT&RUN signal plot is displayed above. **(e)** Venn diagram showing the overlap between genes with CENP-B at promoter at G2 and clustered histone genes. **(f)** Bar chart showing the percentages of clustered histone gene promoters with and without CENP-B containing the specified number of CCAAT boxes. Data were analysed by Fisher’s exact test (** p = 0.0017), comparing count distribution in bar 1 to the average count distribution across bars 2-4 (negative controls). **(g)** Violin plots of MFE predictions for clustered histone gene promoters with and without CENP-B. Data were analysed by two-tailed Welch’s t-test (**** p < 0.0001), the median is marked by a solid line, and quartiles are marked by dashed lines. **(h-i)** Heatmaps showing the normalised transcript per million (TPM) ranks (0 = lowest ranking, 1 = highest ranking) of clustered histone genes in G1-and G2-synchronised RPE1 cells treated with non-targeting siRNA (h), and in unperturbed asynchronous and S phase RPE1 cells (i), showing genes with and without CENP-B at promoter in G2. **(j)** Volcano plots showing expression changes of the clustered histone genes in CENP-B siRNA-treated compared to non-targeting siRNA-treated cells, for G1 and G2-synchronised cells. The transcript log2FC (x-axis) and-log10(p-adj) (y-axis) are shown, genes with CENP-B binding at their promoter are in red. (Horizontal dashed lines indicate p-adj = 0.05)

Histone genes are organised into four clusters in the human genome^23^. Interestingly, two of the significant nc-CENP-B bound clusters correspond to histone gene clusters 1 and 2, located on chr6 and chr1, respectively (Fig. 6a-d). We therefore investigated this relationship further. Of the 85 histone genes from all four histone gene clusters that are expressed in a replication-dependent manner^23^, we found CENP-B is associated with the promoters of approximately half (39 promoters, 46%) of these genes (Fig. 6e). Notably, the CENP-B binding pattern is well conserved in both K562 and HEK293 cell lines (Fig. 6b-d and Supplementary Fig. 9a), suggesting this is functionally important.

Consistent with other nc-CENP-B sites, CENP-B shows stronger histone gene promoter enrichment in G2 than G1 phase cells in all cell lines (Supplementary Fig. 9b-d). Unlike the larger cohort of nc-CENP-B binding sites, however, significant association of other centromere associated proteins is not apparent at the histone gene promoters (Supplementary Fig. 9e).

CENP-B-bound histone gene promoters are more likely to contain multiple CCAAT boxes (Fig. 6f) and these are predicted to form more stable secondary structures (Fig. 6g), suggesting that the determinants for nc-CENP-B binding are shared at these sites. Histone gene regulation is tightly coordinated and reaches a peak in S phase^23^. We therefore interrogated RNA-seq data from asynchronous and S phase RPE1 cells^24^ and found that CENP-B is more likely to bind to highly expressed histone genes, and this is also apparent when we look at expression levels in G1 or G2 phase cells (Fig. 6h,i).

While only a handful of histone genes was significantly misregulated in our RNA-seq dataset, we nevertheless looked in more depth at the impact of CENP-B depletion on histone gene expression since modest mis-regulation across the entire set of histone genes can have important consequences. Strikingly, we find that CENP-B depletion leads to increased expression of almost all histone genes present in our dataset, particularly in G2 phase cells, regardless of whether CENP-B was found at the promoter (Fig. 6j). These data suggesting that CENP-B normally constrains histone gene expression outside of S phase and are consistent with a regional role for CENP-B in regulating expression of gene clusters.

## DISCUSSION

Here, we demonstrate that CENP-B binds to sites in chromosome arms. This association is dependent on the CENP-B DNA binding domain, but not on the presence of B box motifs. Instead, nc-CENP-B binds to sites characterised by the presence of multiple repeated motifs, often CCAAT boxes, that can form secondary structure. These nc-CENP-B sites also coincide with regions of negative supercoiling, which is likely driven by RNApolII activity, given the prevalence of divergently transcribed promoters present in nc-CENP-B bound locations. Such conditions favour secondary structure formation, and in support of this, we find CENP-B binds hairpin-forming DNA in vitro. Notably, however, additional nc-CENP-B binding determinants must exist, given the abundance of both CCAAT boxes and secondary structure forming sequences in the human genome.

Our data demonstrate that gene expression changes when CENP-B is depleted are not obviously linked to the presence of CENP-B on those promoters under unperturbed conditions. This suggests that CENP-B is not functioning as a conventional transcription factor. As CENP-B mediates loop formation in centromeres^25^, a potential model is that nc-CENP-B also contributes to in chromosome arm regions (Supplementary Fig. 9f).

While we found no strong relationship between nc-CENP-B sites and TAD boundaries, suggesting it is not working through this pathway, recently, the acidic region of CENPB was shown to contribute to the reorganisation of chromosomes to facilitate centromere organisation during mitosis^9^. Although the chromatin environment is markedly different, it is possible that this property of CENP-B, together with its ability to form loops^25^, might also influence clustering or 3D organisation of chromosome arm regions. Notably, in this regard, the replication-dependent histone genes normally assemble in the G1/S phases of the cell cycle into a Histone Locus Body (HLB) to facilitate gene expression during S phase, and this structure is dissolved as cells move into G2 phase^26,27^. One attractive possibility is that nc-CENP-B, by virtue of its ability to influence chromatin organisation, coordinates the reorganisation of the HLB to facilitate down-regulation of these genes in G2 phase cells (Supplementary Fig. 9g).

A mechanism along these lines would fit with the lack of direct correlation between promoter binding and impact on gene expression and would support a paradigm in which groups of nc-CENP-B bound loci are assembled into transcription-permissive or-repressive higher-order structures. Loss of CENP-B would therefore have the potential to impact the local environment of groups of genes, in a manner akin to transcription compartments^28^.

Moreover, like transcription compartments, whether the expression of any individual gene in this region is altered by CENP-B-dependent structural organisation will depend on a range of other factors, such as the strength of transcriptional regulators of each gene.

We see a modest association of CENP-A and other CCAN components - CENP-H, CENP-K, CENP-T, and CENP-W - with nc-CENP-B binding sites. Whether this reflects assembly of a ‘mini-CCAN’ or looping such that nc-CENP-B sites transiently encounter centromeres is not yet clear. There is evidence that CENP-A can localise to promoters and drive gene expression, particularly when overexpressed in cancer cells^29^. CENP-B is also often overexpressed in cancer (for example, ^30^), and it will be of interest to determine the interplay between these factors in chromosome arms when misregulated.

These findings provide a new dimension to CENP-B biology, and further investigation will determine whether the non-centromeric functions of CENP-B contribute to pathological contexts, such as cancer, where cell cycle regulation plays a central role.

## MATERIALS AND METHODS

### Cell culture

hTERT-RPE1 were obtained from ATCC (catalogue number CRL-4000) and cultured in Dulbecco Modified Minimal Essential Medium (DMEM)/F-12 (Sigma) supplemented with 10% FBS (Gibco), 200 µM glutamax (Gibco), 0.26% sodium bicarbonate (Gibco), and 1% penicillin/streptomycin (P/S) (Sigma). HEK293TN cells were obtained from ATCC and cultured in DMEM supplemented with 10% FBS (Gibco), and 1% penicillin/streptomycin (P/S) (Sigma). Cells were maintained at 37°C in a humified incubator with 5% CO2 and were regularly tested for mycoplasma contamination.

### CUT&RUN sequencing

CUT&RUN (Cleavage Under Targets & Release Using Nuclease) was performed according to the CUT&RUN Assay Kit protocol (Cell Signaling Technology) with the following modifications. Cells were detached with Accutase (Sigma). 2×10^5^ cells were collected per experiment and pelleted by centrifugation for 5 minutes at 600 x g. Beads were incubated in the indicated primary antibody (Table S1) on a nutator overnight at 4°C. MNase digestion was carried out at 0°C in an ice water bath for 30 minutes. Salt fractionation and DNA purification were carried out according to the CUT&RUN.salt protocol ^22^. Briefly, after STOP buffer addition, samples were incubated at 4°C for 1 hour, and the supernatant containing the low salt fraction was collected. Beads were resuspended in low salt buffer (175 mM NaCl, 10 mM EDTA, 2 mM EGTA, 0.1% TritonX-100, 20 μg/mL glycogen). High salt buffer (825 mM NaCl, 10 mM EDTA, 2 mM EGTA, 0.1% TritonX-100, 20 μg/mL glycogen) was added to beads dropwise with gentle vortexing. Samples were again rocked at 4°C for 1 hour, centrifuged for 5 minutes at 16,000 x g, and the supernatant containing the high salt fraction was collected. Low salt fractions were adjusted to 500 mM NaCl. RNAse A

(Thermo Fisher Scientific) was added to all fractions and samples were incubated at 37°C for 20 minutes. SDS (sodium dodecyl sulfate) and Proteinase K (Cell Signaling Technology) were added and samples were incubated at 50°C for 1 hour. DNA was extracted using phenol/chloroform and precipitated in 100% ethanol as described before library preparation. Libraries were prepared using the NEBNext Ultra II DNA library prep kit for Illumina (New England Biolabs), profiled using the Agilent TapeStation D1000 high sensitivity ScreenTape on the Agilent 4150 TapeStation System, and sequenced on the Illumina Novaseq 6000 with paired-end reads (150 bp in length) by Novogene (Novogene Corporation Inc. UK).

### CUT&RUN data processing

Fastq reads were trimmed using TrimGalore (v0.6.6) (Krueger F, Trimgalore (2023), GitHub repository, https://github.com/FelixKrueger/TrimGalore) using the options --trim-n --paired. The human CHM13-T2Tv1.1 (GenBank GCA_009914755.3) and *S. cerevisiae* S288C (GenBank GCA_000146045.2) assemblies were combined into one FASTA file, then trimmed reads were mapped using Bowtie2 (v2.4.2) ^31^ using the parameters --local --very-sensitive-local --no-unal --no-mixed --no-discordant --dovetail --soft-clipped-unmapped-tlen--non-deterministic --phred33-I 50-X 1500. Reads with more than 3 mismatches were removed with Sambamba (v0.5.0) ^32^, and their corresponding mates were removed with Picard tools (v2.23.8) (“Picard Toolkit.” 2019. Broad Institute, GitHub Repository. https://broadinstitute.github.io/picard/; Broad Institute). Sam files were converted to bam with SAMtools (v1.11) ^33^. Reads mapped to the CHM13-T2Tv1.1 assembly were extracted using BAMtools (v2.5.1) ^34^ split. Low and high salt BAM files from each sample were merged, sorted, and indexed using SAMtools. bigwig files were generated from bam files (generated as described in chapter 2.3.1) using deepTools ^35^ bamCoverage --extendReads -- binSize 1 and scaled according to the total number of mapped reads. Peak calling was performed using MACS2 (v2.2.7.1) ^36^ callpeak using merged low and high salt bam files and IgG as negative control, with options-g 3054832041-f BAMPE --keep-dup all-q 0.01 -- broad --broad-cutoff 0.01. CUT&RUN peaks for each condition and biological replicate were called across the genome against their corresponding IgG negative control, using the MACS2 “broad peak” setting and a statistical significance cutoff of FDR q-value < 0.01. Samples (target and control) were normalised to each other during peak calling by the total number of reads in the input bam file and duplicate reads were retained. Peaks within centromere HORs were filtered out using Bedtools ^37^. Bedtools multiiner was used to identify peaks present in 3 out of 3 replicates. Read coverage tracks and peaks were visualised on IGV ^38^. Heatmaps were created using EnrichedHeatmaps (v1.36.0) ^39^ on R.

For spike-in normalisation and visualisation, the number of reads mapped to the *S. cerevisiae* genome was calculated using SAMtools stats. Spike-in normalised bigwig tracks were generated using deepTools --extendReads --binSize 1 and scaled according to the number of *S. cerevisiae* mapped reads.

The K562 CENP-B CUT&RUN data (GSE104805)^22^ was obtained from SRA, with the 4 CENP-B CUT&RUN replicates (low salt and high salt fractions) sequenced at paired-end 25 bp reads used for analysis. Data was analysed as described above with the following changes. Trimmed reads were mapped to the combined human CHM13-T2Tv1.1 and S. cerevisiae S288C reference genomes using Bowtie2 (v2.4.2)^31^ --end-to-end --no-unal --no-mixed --no-discordant --non-deterministic --phred33-I 10-X 700. Bedtools multiiner was used to identify peaks present in 4 out of 4 replicates. For overexpressing HA-CENP-B and HA-CENP-B^ΔDBD^ CUT&RUN, data processing and analysis was performed as described above with the following changes. Peak calling was performed using MACS2 (v2.2.7.1) callpeak using merged low and high salt bam files and IgG as negative control, with options-g 3054832041-f BAMPE --keep-dup all-q 0.00001 --broad --broad-cutoff 0.00001. CENP-B peaks were filtered for non-centromeric peaks using Bedtools.

### Motif enrichment analysis

Motif enrichment analysis of non-centromere CENP-B peaks was performed using SEA (Simple enrichment analysis of motifs) ^40^ on the MEME suite (v5.4.1) ^41^. Consensus CENP-B peak co-ordinates were converted to FASTA sequences using Bedtools getfasta. Motif enrichment analysis was performed using the human CENP-B motif from the HOCOMOCO (v11) human collection or the full HOCOMOCO (v11) human collection ^42^. The co-ordinates of all CCAAT motifs in the CHM13-T2Tv1.1 reference genome was identified using IGV Find Motif. Bedtools shuffle was used to randomly permute the genomic locations of the nc-CENP-B peaks (excluding centromeric region) or CCAAT boxes. The number of CCAAT boxes within each nc-CENP-B peak was determined using Bedtools intersect-c.

### Promoter analysis

Promoters were defined as the region between 1 kb upstream and 0.2 kb downstream of a gene’s transcription start site (TSS), as defined by GENCODE v35 CAT+ Liftoff annotations in CHM13-T2T. Promoters bound by CENP-B were determined by Bedtools intersect. Gene ontology analysis for the CENP-B-bound gene promoters were performed using ShinyGO ^43^, with the human Hallmark ^44^ and Gene Ontology: Biological Processes ^45^ gene sets, using a statistical significance cut-off of FDR < 0.05.

### Minimum free energy predictions

Peak bed files were converted to fasta using Bedtools getfasta. Minimum free energies (MFE) of peak sequences were predicted using the ViennaRNA ^46^ package, with the command RNAfold-d2-g --noLP-P dna_mathews2004.par --noPS –noconv ^20^. For consensus nc-nc-CENP-B peaks in RPE1, K562, and HEK293, the full set of peaks and their shuffled controls were used in MFE analysis.

For consensus centromeric CENP-B peaks in RPE1, K562, and HEK293, peaks were divided into 1000 bp windows using Bedtools makewindows-w 1000 and keeping peaks above 50 bp wide. 1000 of these windows were sampled for each cell line using Bedtools sample-n 1000. These 1000 peaks were randomised using bedtools shuffle. MFE predictions were performed as described above.

### ATAC-seq

ATAC-seq (Assay for transposase-accessible chromatin by sequencing) was performed according to the Omni-ATAC protocol ^47^. Briefly, 1 x 10^5^ cells were collected per experiment and nuclei were extracted. Nuclei were incubated with Tn5 on ice for 30 minutes. DNA was purified using the Qiagen MinElute reaction cleanup kit (Qiagen), and amplified using the NEBNext high-fidelity PCR master mix (New England Biolabs) and Illumina Nextera DNA indexes (Illumina). Libraries were purified using AMPure XP beads (Beckman Coulter), then profiled using the Agilent TapeStation D1000 high sensitivity ScreenTape on the Agilent 4150 TapeStation System, and sequenced on the Illumina Novaseq 6000 with paired-end reads (150 bp read length) by Novogene (Novogene Corporation Inc. UK).

### ATAC-seq analysis

Fastq reads were trimmed using TrimGalore (v0.6.6) (Krueger F, Trimgalore (2023), GitHub repository, https://github.com/FelixKrueger/TrimGalore) using the options --trim-n --paired. Trimmed reads were mapped to the human CHM13-T2Tv1.1 genome (GenBank GCA_009914755.3) using Bowtie2 (v2.4.2) ^31^ using the parameters --local --very-sensitive-local --no-unal --no-mixed --no-discordant --dovetail --soft-clipped-unmapped-tlen --non-deterministic --phred33-I 50-X 1000. Reads with more than 3 mismatches were removed with Sambamba (v0.5.0) (Tarasov et al., 2015), and their corresponding mates were removed with Picard tools (v2.23.8) (“Picard Toolkit.” 2019. Broad Institute, GitHub Repository. https://broadinstitute.github.io/picard/; Broad Institute). Sam files were converted to bam with SAMtools (v1.11) ^33^. Reads mapped to chrM were removed using SAMtools, and duplicate reads were removed using Picard. Bigwig files were generated using deepTools bamCoverage ^35^. Peaks were called using MACS2 (v2.2.7.1) ^36^ callpeak with filtered bam files as input and options-g 3054832041-f BAMPE-0.01. ATAC-seq peaks for each biological replicate were called across the genome using the MACS2 “narrow peak” option with a statistical significance cutoff of p-value < 0.01. Bedtools ^37^ multiiner was used to identify peaks present in 2 out of 2 biological replicates.

### Divergent transcripts analysis

Divergent transcripts were determined as described in ^48^ using T2T-CHM13v2.0 GENCODE v35 ^49^ gene annotations. TSSs (Transcription start sites) that were within 200 bp of the same ATAC-seq peak on both forward and reverse strands were defined as bidirectional transcripts.

### siRNA-mediated gene depletion

100 µM of the specified CENP-B siRNA pool (Dharmacon) were transfected into cells using Lipofectamine RNAiMAX (Invitrogen), according to the manufacturer’s instructions.

Reverse transfection was performed for G1-synchronised cells and forward transfection was performed for G2-synchronised cells. Pooled non-targeting siRNAs (Dharmacon) siGENOME non-targeting control siRNA pool 2) were used as control.

### Cell cycle synchronisation

Cells were synchronised at G1 using 1 µM Palbociclib (Cayman Chemical) for 18 hours. Cells were synchronised at G2 using 9 µM RO-3306 (Sigma) for 19 hours. For RNA-seq experiments, cell cycle synchronisation was performed simultaneously as siRNA-mediated gene depletion. Synchronisation was confirmed by flow cytometry as detailed below.

### Flow cytometry

For flow cytometric analysis of cell cycle distribution, growth medium was collected, and cells were trypsinised and combined with cells suspended in growth medium. Cells were pelleted by centrifugation and washed once with PBS. After removal of most supernatant, cells were gently resuspended in remaining PBS and fixed by addition of 70% ethanol dropwise with gentle vortexing and incubated at-20°C overnight. Cell suspensions were centrifuged for 5 minutes at 300 x g and the ethanol removed. Pellets were washed in cold PBS and then resuspended in the appropriate volume of PBS containing 5 µg/mL propidium iodide (Sigma) and 0.1 mg/mL RNase A. Cells were incubated at 37°C for 30 minutes, protected from light. DNA content of at least 10,000 cells per condition was detected on a BD LSR II (BD Biosciences). Cell debris and doublets were removed by gating and cell cycle phases were quantified using FlowJo software (v10.10.0).

### RNA extraction and sequencing

For RNA-sequencing, cells were harvested by trypsinisation followed by centrifugation. 3 independent biological replicates were used for G1 cells, and 2 biological replicates were used for G2 cells. RNA was purified using the NEB Monarch total RNA Miniprep kit (New England Biolabs), quantified by Nanodrop (Thermo Scientific), and profiled using the Agilent High Sensitivity RNA ScreenTape, using the Agilent Tapestation as above. Library preparation and sequencing was performed by Novogene Corporation Ltd. Novogene NGS Stranded RNA Library Prep Set was used to generate 250-300 bp insert strand specific libraries, and ribosomal RNA was removed using TruSeq Stranded Total RNA Library Prep, followed by ethanol precipitation. RNA was fragmented to 250-300 bp. After fragmentation, the first strand cDNA was synthesized using random hexamer primers. During the second strand cDNA synthesis, dUTPs were replaced with dTTPs in the reaction buffer. This was followed by end repair, A-tailing, adapter ligation, size selection, USER enzyme digestion, PCR amplification, and final library purification. The libraries were checked with Qubit and real-time PCR for quantification and bioanalyzer for size distribution detection. 50 million paired-end reads (150 bp read length) were sequenced on an Illumina NovaSeq X.

### RNA-sequencing analysis

Fastq reads were checked using FastQC (v0.11.9) (Andrews, S. (2010). FastQC: A Quality Control Tool for High Throughput Sequence Data [Online]. http://www.bioinformatics.babraham.ac.uk/projects/fastqc/) and trimmed using TrimGalore (v0.6.6) (Krueger F, Trimgalore (2023), GitHub repository, https://github.com/FelixKrueger/TrimGalore). Residual ribosomal RNA reads were removed using Ribodetector (v0.2.7) ^50^ with-e norRNA setting and strandedness was detected using RSeQC (v4.0.0) ^51^. Reads were aligned to the T2T-CHM13v2 genome using STAR alignment software (v2.7.6a) ^52^ and reads mapping to genes were quantified using HTSeqCount (v0.12.4) ^53^. Differential analysis of gene expression was calculated in R using DESeq2 (v1.38.3) ^54^. Samples were normalised according to sequencing depth by DEseq2 before differential expression analysis. DEseq2 was used to compare gene expression levels between the siCENPB and non-targeting siRNA-treated cells in G1 and G2. Multiple hypothesis testing was performed by DEseq2 to determine the adjusted p-values (p-adj) for the statistical significance in differential expression of each gene. Genes with a p-adj of < 0.05 were used for further analysis and intersection with the G1 and G2 CENP-B CUT&RUN datasets.

Volcano plots were made in R using EnhancedVolcano (v1.24.0).

### Gene cluster analysis

The CHM13-T2T reference genome was tiled into 2 Mb bins with 0.5 Mb sliding window. The densities of genes misregulated (p < 0.05) in G2, genes with CENPB binding at promoter in G2, and all genes (background) in each bin were calculated. Enrichment ratios for misregulated genes / all genes, and genes with CENP-B binding / all genes were calculated per bin.. Enriched regions were called as regions with enrichment ratios > 2 SD. Genome plots were created using KaryoploteR (1.32.0) ^55^ on R.

### Histone gene expression analysis

RPE1 RNA-seq data for asynchronous and S phase ^24^, G1 and G2 (non-targeting siRNA) were used to determine levels of histone gene expression. Histone genes within the 4 clusters ^56,57^ were included in analysis, where TPMs for these genes only were computed from raw counts (excluding N/As). TPMs for each gene in every sample and replicate were ranked, and the average ranks between biological replicates were calculated. For S phase, data from T2-10 were included, average TPM at each timepoint were calculated, the peak TPM within S phase for each gene were selected and ranked. Heatmaps showing TPM ranks and CENP-B binding at promoter for the selected histone genes in asynchronous, S, G1, and G2 were created in R using ComplexHeatmaps (v2.22.0) ^58^.

### Whole protein extraction

Cells were detached by trypsinisation and pelleted by centrifugation. Cell pellets were resuspended in the appropriate volume of lysis buffer (10% glycerol, 50 mM Tris-HCl (pH 7.4), 0.5% NP-40, 150 mM NaCl) containing 0.25 U/µL benzonase nuclease (Sigma), 1x cOmplete™ EDTA-free protease inhibitor cocktail (Roche), and 1x PhosSTOP™ phosphatase inhibitor cocktail (Roche). Cells were lysed on ice for 45 minutes followed by centrifugation for 15 minutes at 16,000 x g at 4°C. The resulting supernatant containing whole cell protein extracts was collected. Protein concentration was estimated using a Bradford assay (Bio-Rad) according to the manufacturer’s protocol.

### Chromatin fractionation and enrichment

Cell pellets were resuspended in the appropriate amount of nuclear extraction buffer (150 mM Tris-HCl (pH 7.5), 60 mM KCl, 15 mM NaCl, 5 mM MgCl_2_, 1 mM CaCl_2_, 250 mM Sucrose) containing 0.3% NP-40, 1 mM DTT, 1x cOmplete™ EDTA-free protease inhibitor cocktail (Roche), and 1x PhosSTOP™ phosphatase inhibitor cocktail (Roche), then incubated on ice for 5 minutes. Nuclei were pelleted by centrifugation for 5 minutes at 500 x g at 4°C. Nuclear pellet was washed with nuclear extraction buffer with 1 mM DTT, 1x protease inhibitor cocktail, and 1x phosphatase inhibitor cocktail. The pellet was resuspended in hypotonic buffer (10 mM HEPES (pH 7.9), 3 mM EDTA, 0.2 mM EGTA) with 1 mM DTT, 1x protease inhibitor cocktail, and 1x phosphatase inhibitor cocktail, then incubated on ice for 30 minutes to release chromatin. Chromatin fraction was obtained by centrifugation for 5 minutes at 1,700 x g at 4°C. Chromatin fraction was solubilised for SDS-polyacrylamide gel electrophoresis by resuspension in LSB, sonicated with the Bioruptor Pico (Diagenode) for 2 cycles of 30 seconds with a 30 second gap. Samples were centrifuged for 5 minutes at 16,000 x g at 4°C. The chromatin fraction (supernatant) was boiled for 5 minutes at 95°C to denature proteins and run in SDS-PAGE.

### Western blotting

For each western blot, approximately 20 µg of protein extract was combined with LSB containing 25 µM DTT and boiled for 5 minutes at 95°C to denature proteins. Proteins were separated via SDS-polyacrylamide gel electrophoresis and transferred to 0.2 µm nitrocellulose membranes (Fisher Scientific). Successful protein transfer was confirmed by incubating membranes in Ponceau S solution (Sigma) for 5 minutes with rocking and membranes were blocked in 5% milk in TBS buffer containing 0.1% Tween-20 for 1 hour with gentle rocking. Membranes were incubated in the appropriate antibody diluted in blocking buffer at 4°C overnight, washed 3 times with TBS buffer containing 0.1% Tween-20, and incubated in the appropriate horseradish peroxidase (HRP)-conjugated secondary antibody diluted in blocking buffer for 1 hour at room temperature. Proteins were visualised on an iBright CL750 imager (Thermo Fisher Scientific) using Immobilon Crescendo HRP substrate for chemiluminescence.

### CENPB and CENPB^ΔDBD^ protein expression and purification

Both full-length CENPB and CENPB^ΔDBD^ (amino acid 132-599) gene constructs were codon optimized and followed by 3C cleavage site and StrepII^x2^. Both constructs were cloned into pFastBac-1. Bacmids were obtained using DH10 EmBacY cells. P1, P2, and P3 baculoviruses were generated using Sf9 cells. For protein expression, 500 ml of High Five cells at density of 1.5 × 10^6^ cells ml^−1^ were infected with 5 ml of Sf9 cells pre-infected with P2 baculoviruses. High Five cells were pelleted when viability reached 80-90%, approximately 72 hours post infection. Cells were stored at-80°C until processing.

High Five cell pellets expressing CENPB or CENPB^ΔDBD^ were resuspended in lysis buffer containing 50 mM HEPES (pH 8), 300 mM NaCl, 0.5 mM TCEP, 5% Glycerol, 2.5 mM Benzamidine, cOmplete Mini Protease Inhibitor tablet (Roche, 2 tablets for 50 ml lysis buffer), and Benzonase Nuclease (Millipore, 5 ul for 50 ml lysis buffer). Cells were lysed by sonication and insoluble matters were removed by centrifugation. The filtered soluble fraction was subsequently loaded into 5 ml Strep-Tactin Superflow column (Qiagen), pre-equilibrated in 50 mM HEPES (pH 8), 300 mM NaCl, 0.5 mM TCEP, 5% Glycerol. Bound proteins were washed with buffer containing 50 mM HEPES (pH 8), 1000 mM NaCl, 0.5 mM TCEP, 5% Glycerol. Proteins were eluted from the Strep-Tactin Superflow column in buffer containing 50 mM HEPES (pH 8), 300 mM NaCl, 0.5 mM TCEP, 5% Glycerol, 3 mM Desthiobiotin. StrepII^x2^ tag was removed by incubating the protein with 3C protease overnight at 4°C. Final polishing was achieved by size exclusion chromatography using HiLoad 16/600 Superose 200 pg (Cytiva) in buffer containing 20 mM HEPES (pH 8), 150 mM NaCl, 0.5 mM TCEP. Peak fractions were pooled and concentrated up to 20-30 μM.

### CENP-B oligo pull-down assay

HPLC-purified DNA oligos were obtained from IDT, with 3’ Biotin-TEG modification on each forward strand oligo. Oligos were resuspended in 1 x TE (pH 7.5). Oligos were annealed at a concentration of 20 μM in 10 mM Tris HCl (pH 7.4), 100 mM KCl. ssDNA oligos were heated at 95°C for 5 minutes then cooled on ice immediately. dsDNA oligos were heated at 95°C for 5 minutes then gradually cooled at-0.3°C per minute.

For each pull-down, 50 μL of Streptavidin MegneSphere Paramagnetic beads were added to protein LoBind tubes, and washed twice with 300 μL of pull-down buffer (25 mM HEPES pH 7.5, 120 mM KCl, 1 mM MgCl2, 3% BSA, 20% Glycerol, 0.1% Igepal C-630, 0.4 μg/mL salmon sperm DNA, 1 mM DTT). 200 pmol of oligos were added to the washed beads, and incubated with rolling at room temperature for 30 minutes. Beads were washed twice with pull-down buffer. 1.4 pmol of recombinant CENP-B or CENP-B^ΔDBD^ was added to each reaction, then incubated with rolling at 4°C for 1 hour. Beads were washed three times with wash buffer (25 mM HEPES pH 7.5, 120 mM KCl, 1 mM MgCl2, 20% Glycerol, 0.1% Igepal C-630, 1 mM DTT). Beads were resuspended in 1 x LDS buffer with 25 mM DTT. Samples were boiled for 10 minutes, and the supernatant for each reaction were used for Western Blotting following the protocol mentioned above. For input control lanes, 0.1 x of recombinant CENP-B was loaded.

### Generation of RPE1 CENP-B and CENP-B^ΔDBD^ overexpressing RPE1 cells

Plasmids pLV[Exp]-Puro-CMV>HA/hCENPB(1-599):P2A:NLS-EGFP (CENP-B FL) and pLV[Exp]-Puro-CMV>HA/hCENPB(132-599):P2A:NLS-EGFP (CENP-B ΔDBD) were obtained from VectorBuilder. Plasmids were midiprepped using the Qiagen Plasmid Plus kit according to manufacturer’s protocol.

For lentivirus production in HEK293TN cells, 10 μg of plasmid was transfected in combination with the lentiviral packaging and envelope plasmids, pMD2G and psPAX2 using Lipofectamine 3000 (Thermo Fisher Scientific). Viral particle-containing medium was harvested at 48 h and 72 h post transfection, combined and filtered through 0.45 μm filters.

For transduction, viral particle-containing medium was added to RPE1 cells in 10 cm dishes with 6 μg/mL polybrene (Merck) and incubated for 24 h. GFP-positive cells were sorted by the BD FACSymphony S6 Cell Sorter (BD Biosciences) and expanded. CENP-B overexpression was confirmed by Western Blot using Anti CENP-B and anti HA antibodies.

### Software & statistical analyses

The number of biological replicates for each experiment and the statistical analyses used are indicated in the figure legend for each experiment. Graphs were generated and statistical analyses were performed using GraphPad Prism (v10.6.1). Omics data were visualised using the indicated packages in R (v4.4.3) and RStudio (v2026.01.1+403). Figures were generated using Inkscape (v1.4.3). The schematics in Figure 4 were generated using BioRender (biorender.com).

## Data and code availability

Data for CENP-B, H3K27me3 and IgG CUT&RUN in asynchronous RPE1 cells are available on GEO (GSE235294) ^59^. RPE1 ATAC-seq data are available on GEO (GSE308091). CENP-A, CENP-B, CENP-C and IgG CUT&RUN data in K562 cells were obtained from GEO (GSE104805) ^22^. Asynchronous and S phase RPE1 RNA-seq data were obtained from GEO (GSE137448) ^24^. CUT&RUN data generated in this study are available on GEO (GSE324377). RNAseq data generated in this study are available on GEO (GSE324378).

## Acknowledgements

We thank the Institute of Cancer Research Flow Cytometry Facility for support, and we thank the ICR Scientific Computing Team for HPC services. We thank Professors Jon Pines (ICR) and William Earnshaw (University of Edinburgh) for helpful advice and discussion. We thank all members of the Downs Lab for feedback and discussion.

## Author contributions

L.W., K.A.L., and J.A.D. conceived the study and designed experiments. L.W., K.A.L., R.M., A.H., C.N. performed and analysed experiments, analysed data, and created models. H.W. and A.M. provided materials. A.M., N.G., C.A., and J.A.D. acquired funding and provided supervision. L.W. and J.A.D. wrote the manuscript with input from all authors.

## Competing interests

The authors declare no competing interests.

## Notes

### Competing Interest Statement

The authors have declared no competing interest.

### Funding

- Cancer Research UK: DRCRPG-June24/100004
- Medical Research Council: MR/W001276/1, MC_UU_00007/13, MC_UU_00035/6
- Wellcome Trust: 302297/Z/23/Z, 223097/Z/21

