## Supplementary Data for "CENP-B binds hairpin motifs in chromosome arms influencing gene expression"

### Supplementary Data Figure 1

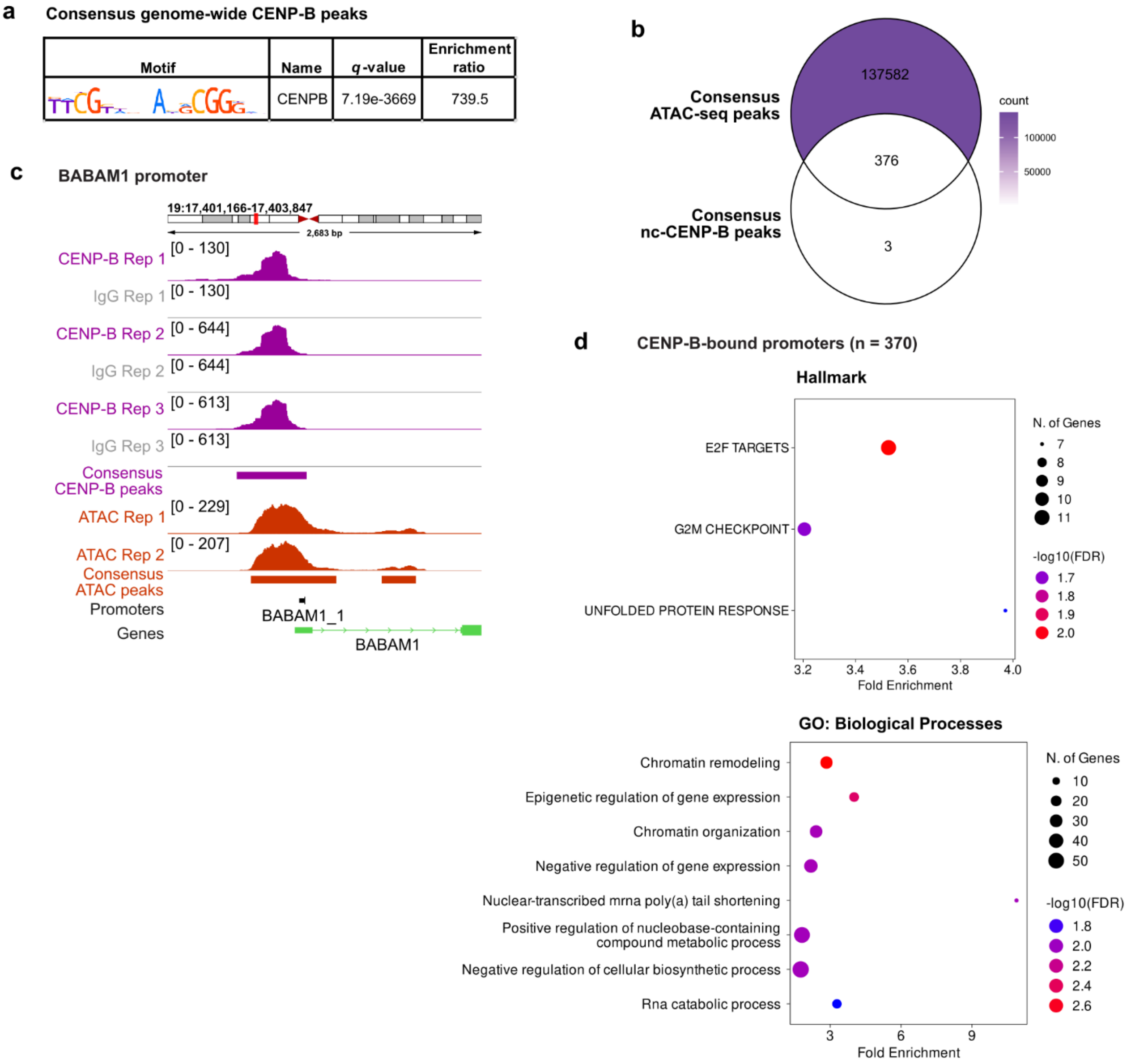


**Supplementary Data Figure 1: CENP-B binding at non-centromere sites**

**(a)** Motif enrichment analysis results for genome-wide CENP-B CUT&RUN peaks in 3 out of 3 replicates. The enrichment ratio indicates the relative enrichment of the CENP-B box motif in the DNA sequences of CENP-B peaks compared to a set of shuffled input sequences. **(b)** Venn diagram showing the number of overlaps between consensus nc-CENP-B peaks (in 3 of 3 biological replicates) and consensus ATAC-seq peaks (in 2 of 2 biological replicates). The colour intensity corresponds to the total number of peaks in each region of the Venn diagram (count). **(c)** IGV screenshot showing the CENP-B binding site at the BABAM1 gene promoter, showing coverage of reads from CENP-B (purple) and IgG (grey) CUT&RUN. ATAC-seq read coverage in RPE1 cells are shown in orange. The locations of the BABAM1 promoter and transcript are shown below. **(d)** Gene ontology (GO) analysis of the Hallmark and GO:Biological Processes sets for the gene promoters (n = 370 genes) occupied by CENP-B in 3 out of 3 biological replicates. The -log(FDR) is shown on the x-axis. The size of the dots represents the number of CENP-B-associated genes in each gene set, and the colour of the dots represents the fold enrichment of the CENP-B-associated genes in the gene set over that of a background gene list.

### Supplementary Data Figure 2

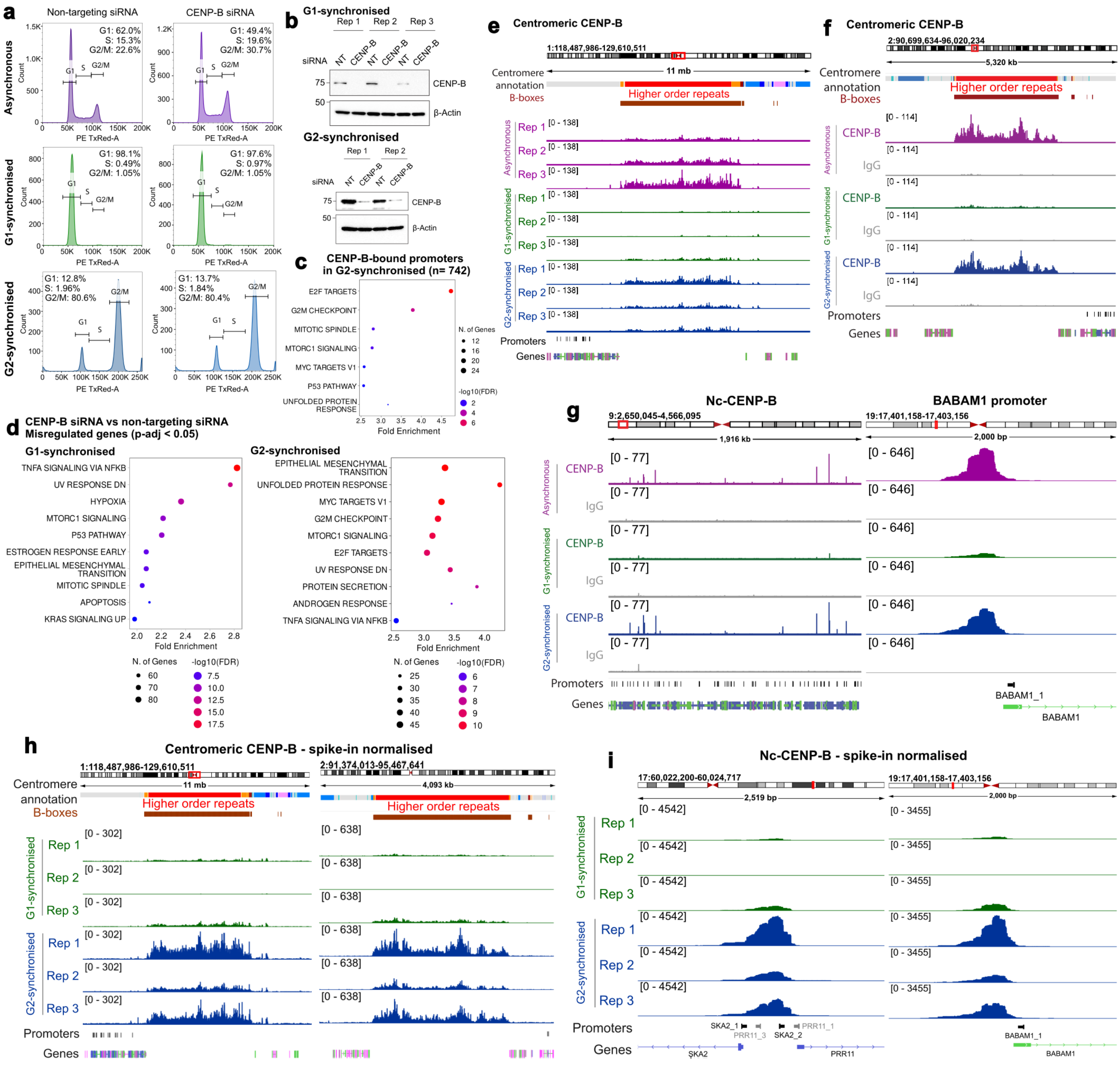


**Supplementary Data Figure 2: CENP-B influences gene expression and shows cell cycle-dependent binding patterns**

**(a)** Histograms showing cell cycle profiles determined by FACS for asynchronous, G1-synchronised, and G2-synchronised RPE1 cells with non-targeting and CENP-B siRNAs. The percentages of cells in G1, S, and G2/M are shown. **(b)** Western blots showing CENP-B protein expression in G1-synchronised RPE1 cells after 18h treatment of Palbociclib, and G2-synchronised RPE1 cells after 19h treatment of RO-3306, treated with non-targeting and CENP-B siRNAs. **(c)** GO analysis of the Hallmark gene sets for promoters with CENP-B binding in G2-synchronised cells. **(d)** GO analysis of the Hallmark gene sets for misregulated genes (p-adj < 0.05) after CENP-B depletion in G1- and G2-synchronised cells. **(e)** IGV screenshot showing coverage of 3 biological replicates of CENP-B CUT&RUN sequencing reads in asynchronous (purple), G1-synchronised (green), G2-synchronised (blue) RPE1 cells across the centromere of chromosome 1. **(f-g)** IGV screenshots showing coverage of CENP-B and IgG CUT&RUN sequencing reads of a representative replicate in asynchronous (purple), G1-synchronised (green), G2-synchronised (blue) RPE1 cells across the centromere of chromosome 2 (f), within the arm of chromosome 9 and at the SKA2/PRR11 promoters (g). For d-f, the centromere HORs are in red, and canonical B-box motifs are in brown. **(h-i)** IGV screenshots showing coverage of spike-in normalised CENP-B CUT&RUN sequencing reads in G1-synchronised (green) and G2-synchronised (blue) RPE1 cells across the centromere of chromosome 1 and 2 (h), and at the SKA2/PRR11 and BABAM1 promoters (i).

### Supplementary Data Figure 3

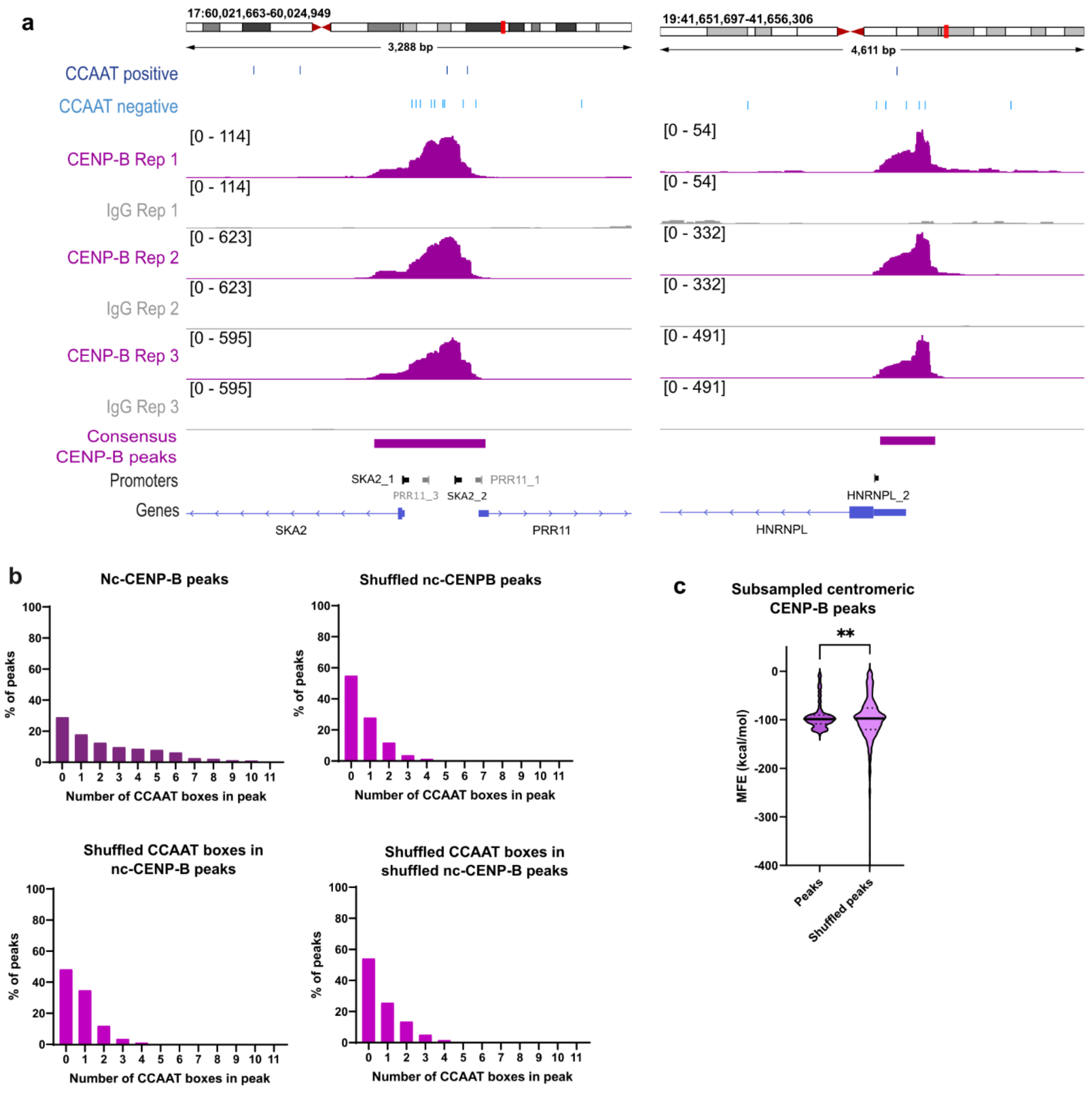


**Supplementary Data Figure 3: Nc-CENP-B peaks are enriched for CCAAT box sequences**

**(a)** IGV screenshots showing CENP-B binding sites at the SKA2 and PRR11 and HNRNPL gene promoters, showing coverage of reads from CENP-B (purple) and IgG (grey) CUT&RUN sequencing in 3 biological replicates. The locations of CCAAT boxes in the reference genome in its positive (dark blue) and negative (light blue) strands are shown above. The locations of promoters and transcripts are shown below. **(b)** Bar charts showing the percentages of CENP-B peaks (y-axis) containing the specified number of CCAAT boxes (x-axis), for nc-CENP-B peaks, shuffled nc-CENP-B peaks, nc-CENP-B peaks with shuffled

CCAAT box co-ordinates, shuffled nc-CENP-B peaks with shuffled CCAAT box co-ordinates. **(c)** Violin plots of MFE predictions for subsampled RPE1 consensus centromeric CENP-B peaks (1000 peaks), and shuffled centromeric CENP-B peaks. Data were analysed by two-tailed Welch's t-test (** $p = 0.0018$), the median is marked by a solid line, and quartiles are marked by dashed lines.

### Supplementary Data Figure 4

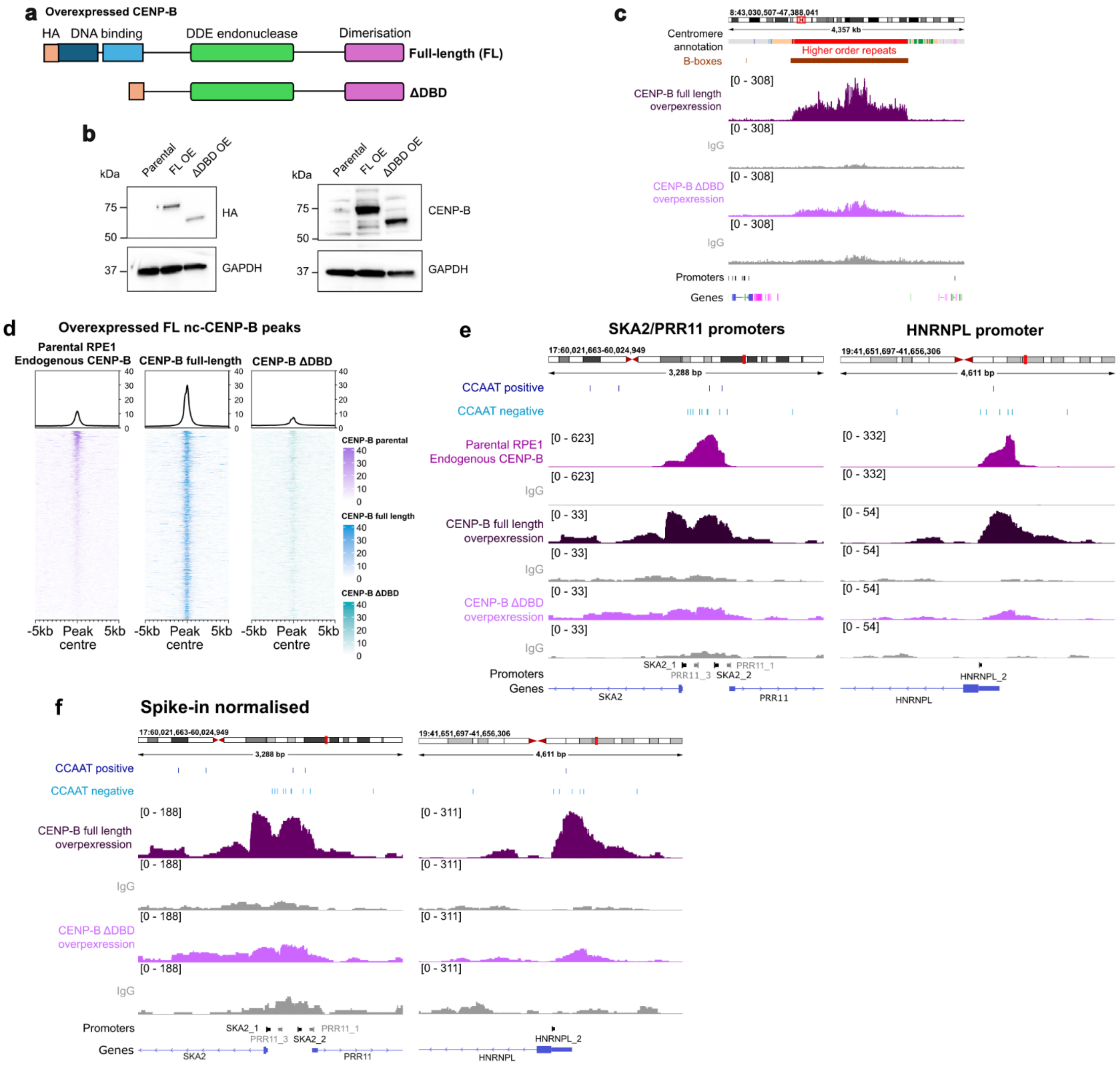


**Supplementary Data Figure 4: CENP-B is directed to non-centromeric sites via its DNA binding domain**

**(a)** Schematics for CENP-B full length (FL) and CENP-B ΔDBD overexpression constructs in RPE1 cells. **(b)** Western blot showing levels of HA and CENP-B in RPE1 parental, CENP-B FL overexpressing, and CENP-B ΔDBD overexpressing cells. **(c)** IGV screenshots showing coverage of CENP-B CUT&RUN sequencing reads for endogenous CENP-B in parental cells (purple), overexpressed CENP-B FL (dark purple) and overexpressed CENP-B ΔDBD (light purple) across the centromere. Active centromere higher order repeats are shown in red, and B-boxes are shown in brown. **(d)** Heatmaps showing CENP-B CUT&RUN

signal for endogenous CENP-B in parental cells (purple), overexpressed full-length CENP-B (dark blue), and overexpressed CENP-B ΔDBD (light blue) at the non-centromere CENP-B peaks bound by overexpressed full-length CENP-B, showing regions +/- 5 kb of each CENP-B peak. An average CUT&RUN signal plot is displayed above each heatmap. **(e)** IGV screenshots showing coverage of CENP-B and IgG CUT&RUN sequencing reads for endogenous CENP-B in parental cells (purple), overexpressed CENP-B FL (dark purple), overexpressed CENP-B ΔDBD (light purple) at the SKA2/PRR11 and HNRNPL gene promoters. The locations of CCAAT boxes are shown above. The locations of promoters and genes are shown below. **(f)** IGV screenshots showing coverage of spike-in normalised CENP-B and IgG CUT&RUN sequencing reads for overexpressed CENP-B FL (dark purple) and overexpressed CENP-B ΔDBD (light purple) at the SKA2/PRR11 and HNRNPL gene promoters.

### Supplementary Data Figure 5

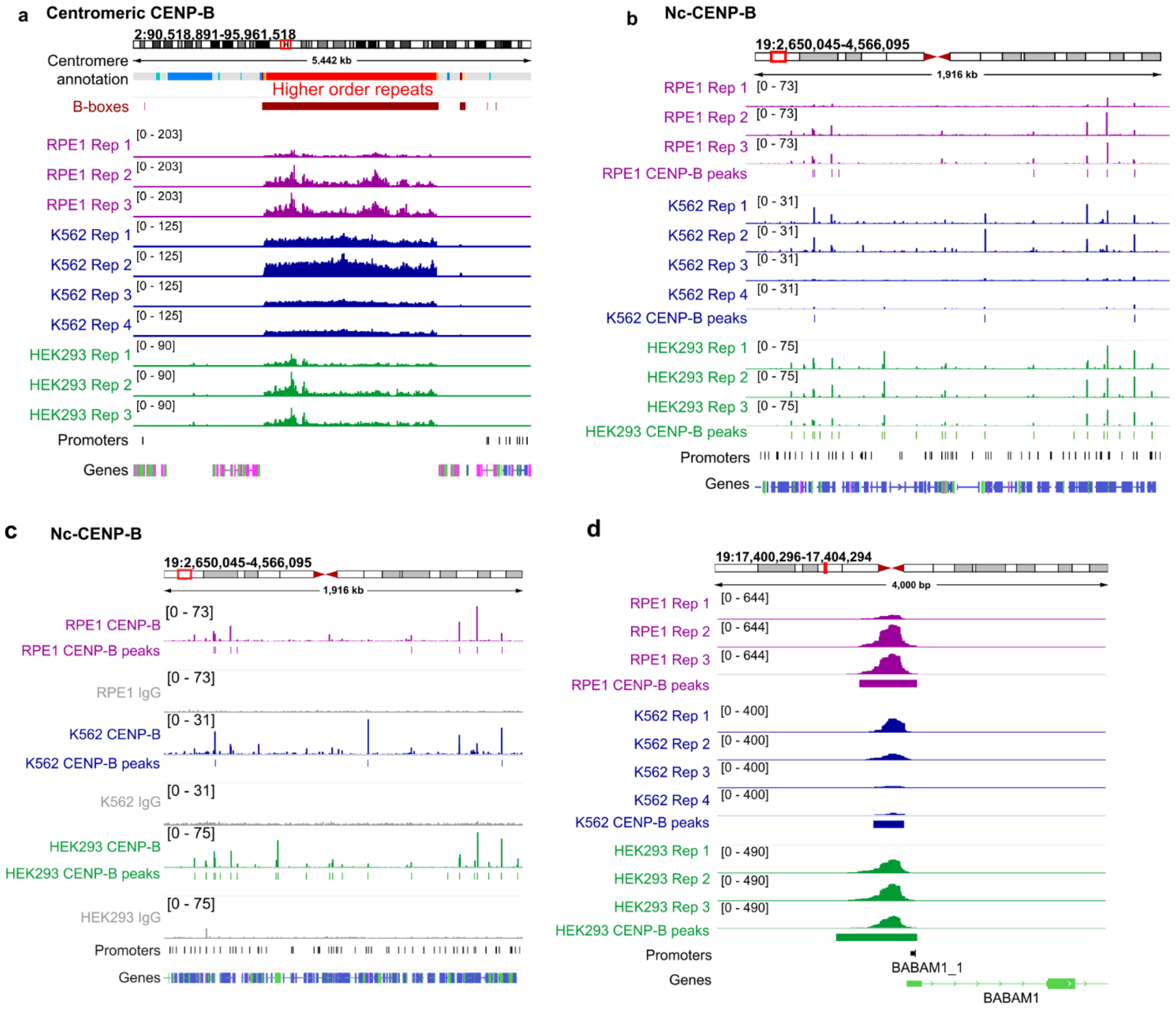


**Supplementary Data Figure 5: Nc-CENP-B binding is conserved between cell types**

**(a-d)** IGV screenshots showing CENP-B binding across the centromere of chromosome 2 (a), within the arm of chromosome 19 (b-c), and at the BABAM1 promoter (d), showing coverage of reads from RPE1 (purple), K562 (blue), HEK293 (green) CENP-B CUT&RUN. The active centromere HORs are shown in red, and canonical B-box motifs are shown in brown. Locations of genes and promoters are shown below.

### Supplementary Data Figure 6

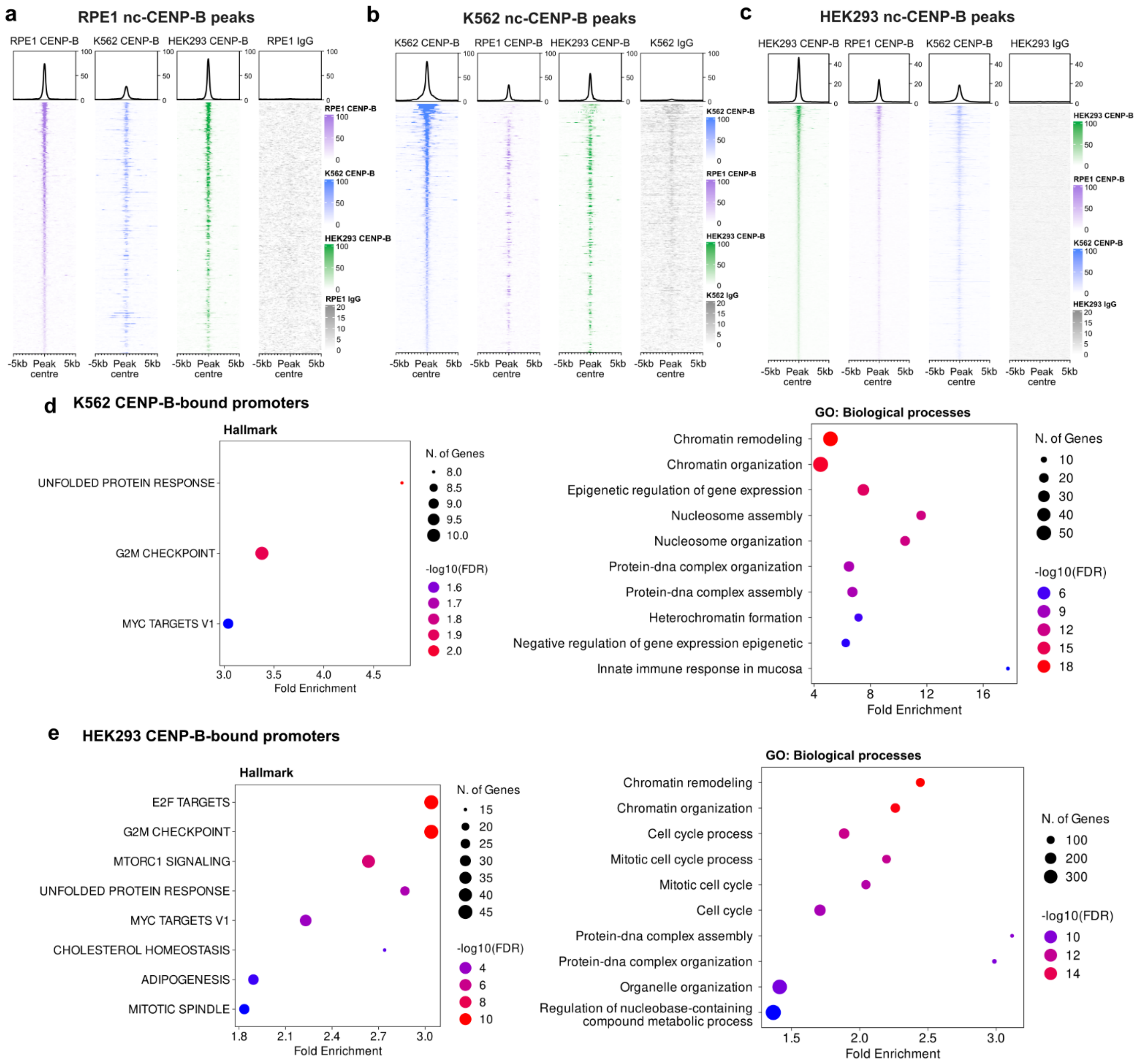


**Supplementary Data Figure 6: Cell type specificity and promoter binding patterns in nc-CENP-B binding sites**

**(a-c)** Heatmaps of CENP-B CUT&RUN signal intensity in RPE1, K562, and HEK293 cells, across the consensus nc-CENP-B peaks in RPE1 (a), K562 (b), and HEK293 (c) cells. An average CUT&RUN signal plot is displayed above. **(d-e)** GO analysis of the top 10 human Hallmark gene sets and GO Biological Processes gene sets for the gene promoters occupied by CENP-B in K562 (d) and HEK293 (e) cells. The -log(FDR) of the gene set enrichment analysis is shown on the x-axis. The size of the dots represents the number of genes within the set of CENP-B-associated genes that are in each gene set, and the colour of the dots represents the fold enrichment of the CENP-B-associated promoters.

### Supplementary Data Figure 7

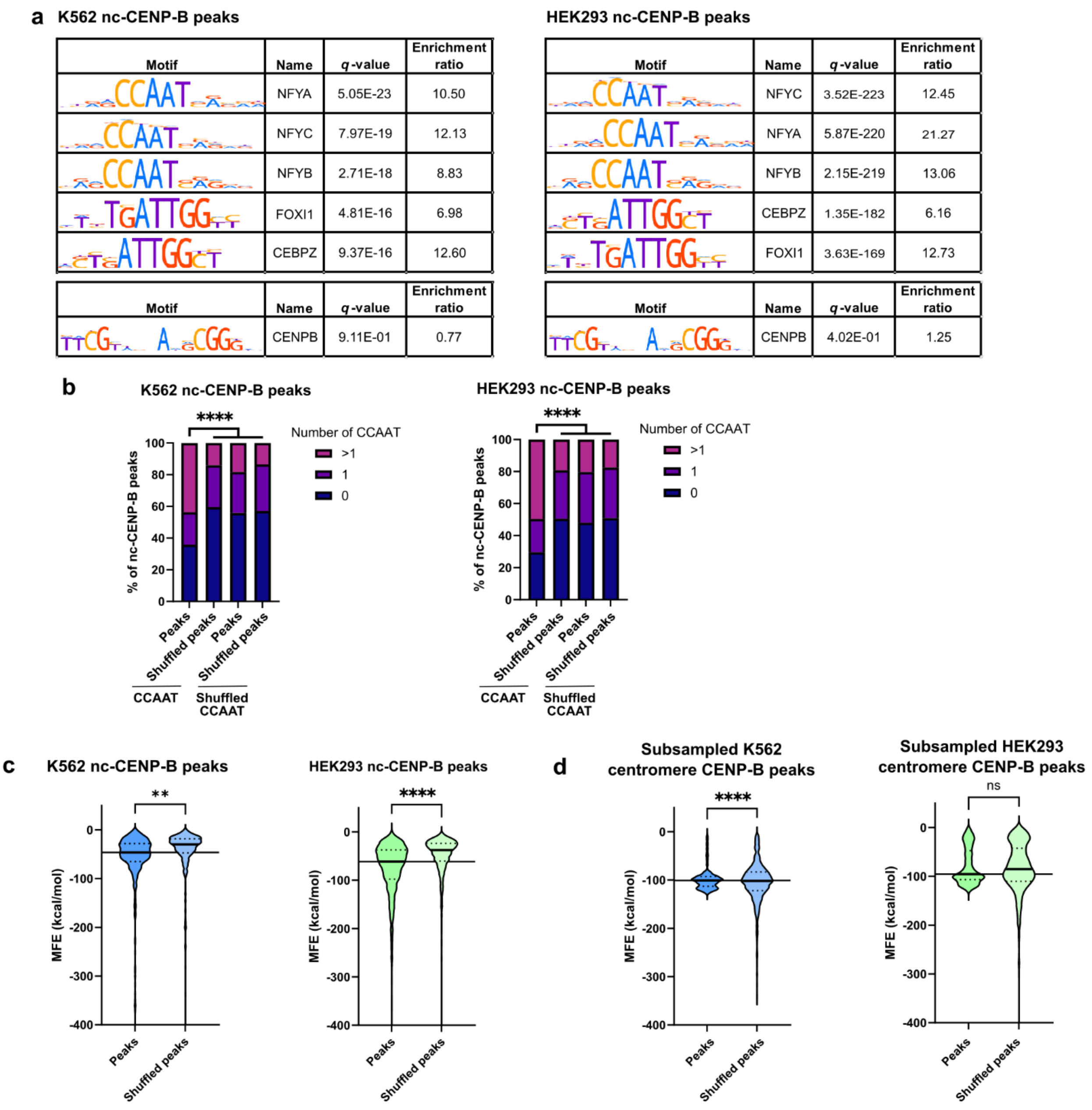


**Supplementary Data Figure 7: Nc-CENP-B binding sites are also enriched for CCAAT boxes in K562 and HEK293 cells**

**(a)** Motif enrichment analysis results for consensus nc-CENP-B CUT&RUN peaks in K562 and HEK293 cells, showing enrichment of the CCAAT box-containing motifs (top) and of the CENP-B box motif (bottom). The enrichment ratio indicates the relative enrichment of each motif in the DNA sequences of nc-CENP-B peaks compared to a set of shuffled input sequences. **(b)** Bar charts showing the percentages of K562 and HEK293 CENP-B peaks containing the specified number of CCAAT boxes and shuffled CCAAT box coordinates for nc-CENP-B peak and shuffled nc-CENP-B peaks. Data were analysed by Chi-square test

(**** p < 0.0001), comparing count distribution in bar 1 to the average count distribution across bars 2-4 (negative controls). **(c)** Violin plots of MFE predictions for K562 and HEK293 consensus nc-CENP-B peaks, and shuffled nc-CENP-B peaks. **(d)** Violin plots of MFE predictions for K562 and HEK293 subsampled consensus centromeric CENP-B peaks (n = 1000), and shuffled centromeric CENP-B peaks. For c-d, data were analysed by two-tailed Welch's t-test test (**** p < 0.0001, ** p = 0.0010, ns = not significant), the median is marked by a solid line, and quartiles are marked by dashed lines.

**Supplementary Data Figure 8**

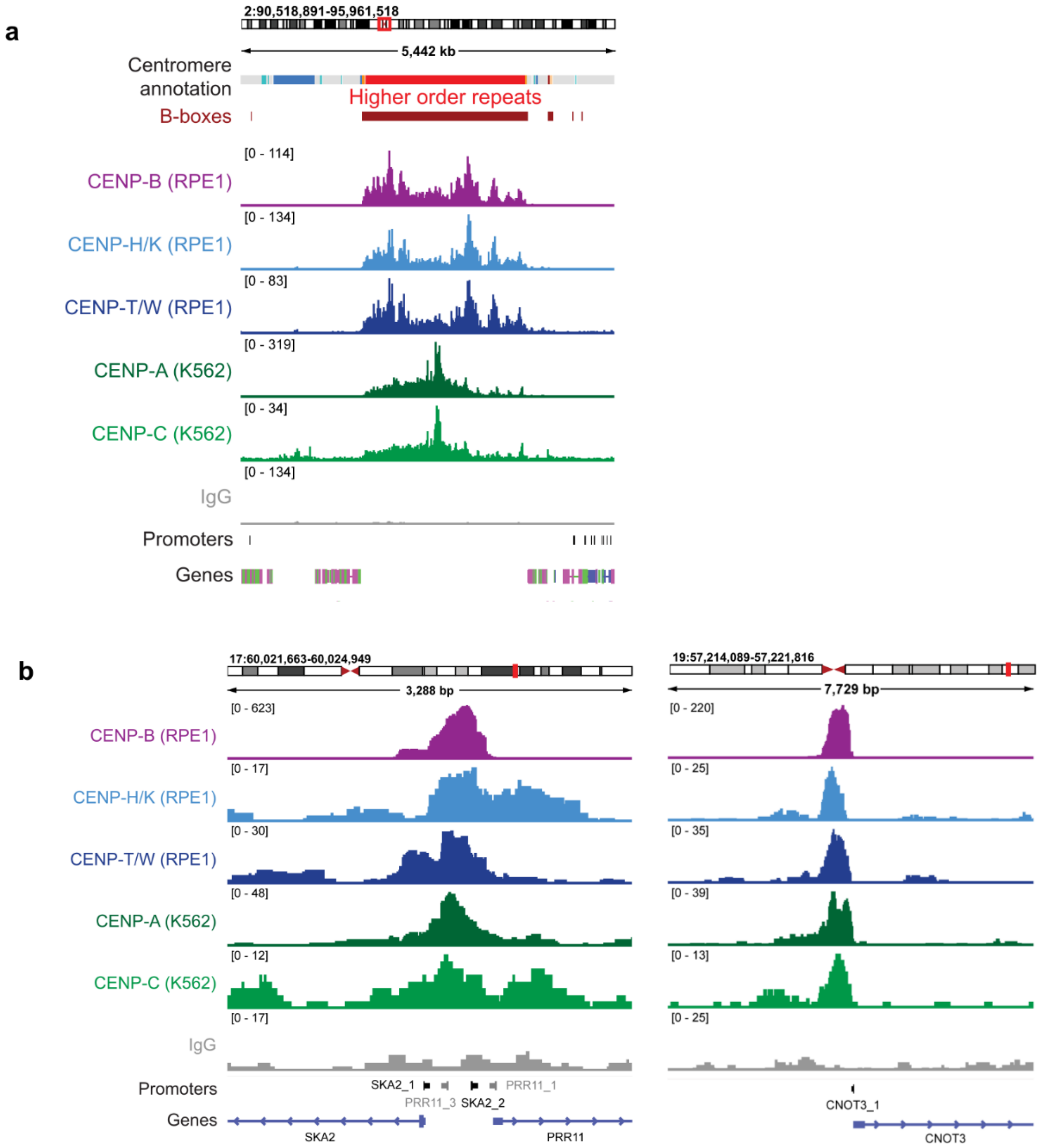


**Supplementary Data Figure 8: Other centromere proteins co-localise with CENP-B at nc-CENP-B sites**

**(a)** IGV screenshot showing enrichment of CENP-B (RPE1), CENP-H/K (RPE1), CENP-T/W (RPE1), CENP-A (K562), and CENP-C (K562) across the centromere of chromosome 2, showing coverage of reads from CENP-B (purple), CENP-H/K (light blue), CENP-T/W (dark blue), CENP-A (dark green), CENP-C (light green), and IgG (grey) CUT&RUN sequencing across the entire centromere. The centromere HORs are shown in red, and canonical B-box motifs are shown in brown. **(b)** IGV screenshots showing CUT&RUN read coverage of CENP-B in RPE1 (purple), CENP-H/K in RPE1 (light blue), CENP-T/W in RPE1 (dark blue), CENP-A in K562 (dark green), CENP-C in K562 (light green), and IgG in

RPE1 (grey) at the SKA2/PRR11 and CNOT3 promoter regions. The locations of the SKA2/PRR11 and CNOT3 promoters and transcripts are shown below each screenshot.

### Supplementary Data Figure 9

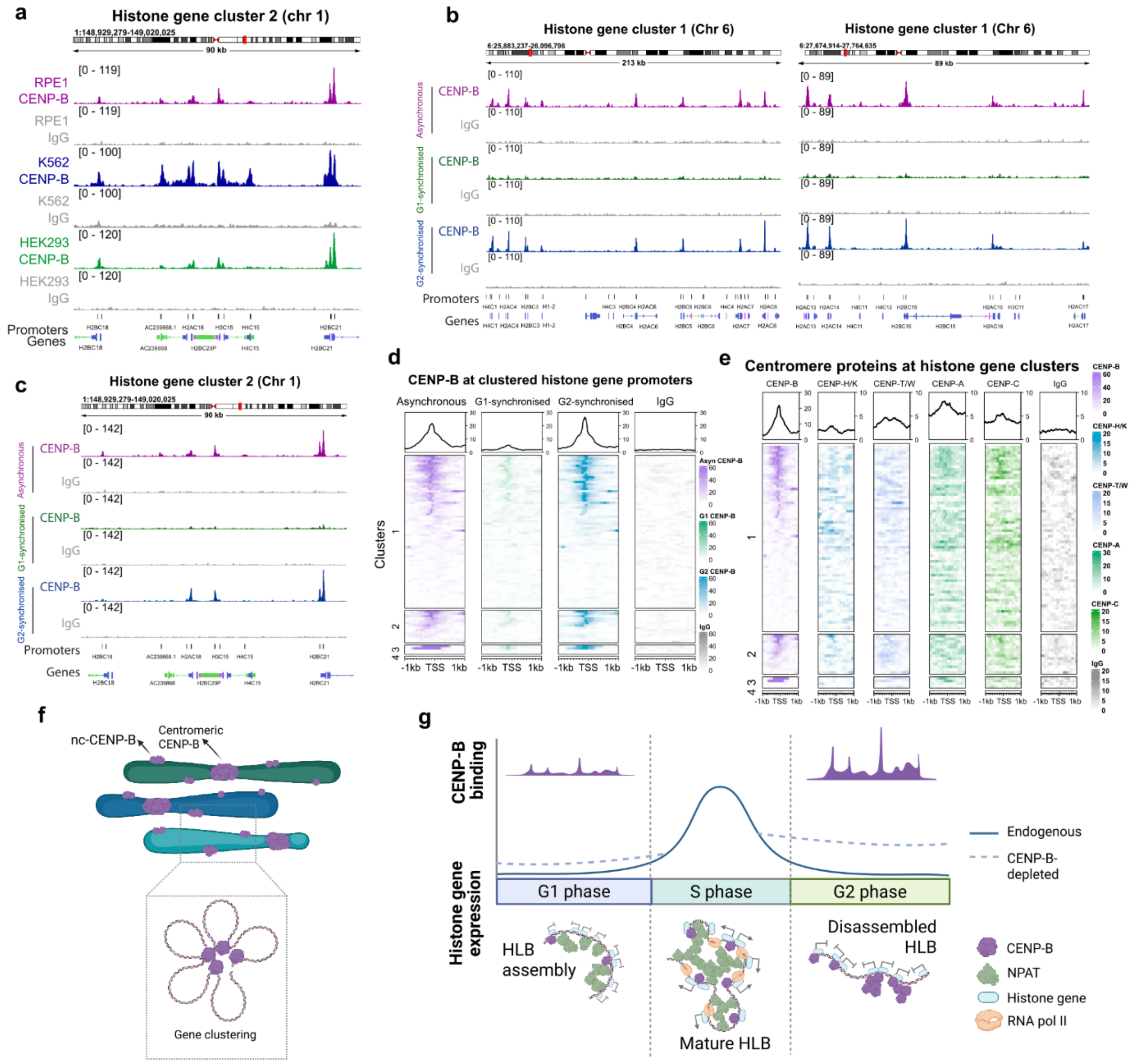


**Supplementary Data Figure 9: CENP-B binds clustered histone gene promoters across cell types**

**(a)** IGV screenshots showing coverage of CENP-B and IgG CUT&RUN reads in RPE1 (purple), K562 (blue), and HEK293 (green) cells at histone gene cluster 2 in chromosome 1. **(b)-(c)** IGV screenshot showing coverage of CENP-B and IgG CUT&RUN reads in asynchronous (purple), G1-synchronised (green), and G2-synchronised (blue) RPE1 cells at (b) two regions within histone gene cluster 1 in chromosome 6 and (c) histone gene cluster 2 in chromosome 1. **(d)** Heatmaps of CENP-B CUT&RUN signal intensity in asynchronous (purple), G1-synchronised (green), and G2-synchronised (blue) cells +/- 1 kb from the TSS of the histone genes within the 4 histone gene clusters. An average CUT&RUN signal plot is displayed above. **(e)** Heatmaps of CUT&RUN signal intensity for CENP-B in RPE1 (purple), CENP-H/K in RPE1 (light blue), CENP-T/W in RPE1 (dark blue), CENP-A in K562 (dark green), CENP-C in K562 (light green), IgG in RPE1 (grey) +/- 1 kb from the TSSs of the clustered histone genes. An average CUT&RUN signal plot is displayed above each heatmap.

All data are from asynchronous cells. **(f)** Model for CENP-B mediating gene clustering across the genome. **(g)** Model for CENP-B function at histone locus bodies (HLBs) at different cell cycle phases. NPAT is a representative HLB-associated protein for illustration.

**Table S1: Antibodies used in this study**

| Target | Source | Catalogue number | Species | Application | Dilution |
|---|---|---|---|---|---|
| CENP-B | Abcam | ab25734 | Rabbit | CUT&RUN | 1:50 |
| | | | | WB | 1:1000 |
| CENP-H/K | Musacchio lab | / | Rabbit | CUT&RUN | 1:50 |
| CENP-T/W | Musacchio lab | / | Rabbit | CUT&RUN | 1:50 |
| H3K27me3 | Cell Signalling Technology | 9733 | Rabbit | CUT&RUN | 1:50 |
| HA | Abcam | ab18181 | Mouse | CUT&RUN | 1:50 |
| | | | | WB | 1:1000 |
| IgG isotype control | Cell Signalling Technology | 66362 | Rabbit | CUT&RUN | 1:20 |
| Actin | Sigma | A5060 | Rabbit | WB | 1:5000 |
| GAPDH | Abcam | ab9485 | Rabbit | WB | 1:2500 |
| Histone H3 | Abcam | ab1791 | Rabbit | WB | 1:5000 |
| Anti-rabbit HRP-conjugated | Agilent Dako | P0448 | Goat | WB | 1:10000 |
| Anti-mouse HRP-conjugated | Agilent Dako | P0260 | Rabbit | WB | 1:10000 |

**Table S2: Oligonucleotides used in this study**

| Name | Sequence (5' – 3') | Application |
|---|---|---|
| siCENPB_1 | GCACGAUCCUGAAGAACAA | CENP-B knockdown |
| siCENPB_2 | GGAGGAGGGUGAUGUUGAU | CENP-B knockdown |
| siCENPB_3 | CCGAAUGGCUGCAGAGUCU | CENP-B knockdown |
| siCENPB_4 | CCAACAAGCUGUCUCCCUA | CENP-B knockdown |
| siControl_1 | UAAGGCUAUGAAGAGAUAC | Non-targeting siRNA |
| siControl_2 | AUGUAUUGGCCUGUAUUAG | Non-targeting siRNA |
| siControl_3 | AUGAACGUGAAUUGCUCAA | Non-targeting siRNA |
| siControl_4 | UGGUUUACAUGUCGACUAA | Non-targeting siRNA |
| B-box_F | TCAGTCAGTCAGTCAGTCAGTCGAAGCAACCT TTGCCCTAGTCAGTCAGTCAGTCAGTCA /3BioTEG/ | CENP-B pull-down |
| B-box_R | TGACTGACTGACTGACTGACTAGGGCAAAGGT TGCTTCGACTGACTGACTGACTGACTGA | CENP-B pull-down |
| Shuffled_B-box_F | TCAGTCAGTCAGTCAGTCAGTCCGCGGTCTACT CTAACAAGTCAGTCAGTCAGTCAGTCA /3BioTEG/ | CENP-B pull-down |
| Shuffled_B-box_R | TGACTGACTGACTGACTGACTTGTTAGAGTAG ACCGCGGACTGACTGACTGACTGACTGA | CENP-B pull-down |
| 2_CCAAT_F | GTCAGTCAGTCAGTCAGTCAGTCCCAATAGTC ATTGGAGTCAGTCAGTCAGTCAGTCAGT /3BioTEG/ | CENP-B pull-down |
| 2_CCAAT_R | ACTGACTGACTGACTGACTGACTCCAATGACT ATTGGGACTGACTGACTGACTGACTGAC | CENP-B pull-down |
| 4_CCAAT_F | AGTCAGTCAGTCAGCCAATAGTCATTGGAGTC ATTGGAGTCCCAATAGTCAGTCAGTCAG /3BioTEG/ | CENP-B pull-down |
| 4_CCAAT_R | CTGACTGACTGACTATTGGGACTCCAATGACTC CAATGACTATTGGCTGACTGACTGACT | CENP-B pull-down |
| 4_CCAAT_2_F | AGTCAGCCAATAGTCATTGGAGTCAGTCAGTC AGTCAGTCATTGGAGTCCCAATAGTCAG /3BioTEG/ | CENP-B pull-down |
| 4_CCAAT_2_R | CTGACTATTGGGACTCCAATGACTGACTGACTG ACTGACTCCAATGACTATTGGCTGACT | CENP-B pull-down |
| 4_CCAAT_3_F | AGTCAGCCAATAGTCAGTCATTGGAGTCAGTC AGTCATTGGAGTCAGTCCCAATAGTCAG /3BioTEG/ | CENP-B pull-down |
| 4_CCAAT_3_R | CTGACTATTGGGACTGACTCCAATGACTGACTG ACTCCAATGACTGACTATTGGCTGACT | CENP-B pull-down |
| No_motif_F | AGTCAGTCAGTCAGTCAGTCAGTCAGTCAGTC AGTCAGTCAGTCAGTCAGTCAGTCAGTC /3BioTEG/ | CENP-B pull-down |
| No_motif_R | GACTGACTGACTGACTGACTGACTGACTGACT GACTGACTGACTGACTGACTGACTGACT | CENP-B pull-down |
| Cen_hairpin_F | CAGATAAAAACTAGAAAGAAGCTTTCTGAGAA ACTTCTTTGTGTGTTACACCTTTCTTTT /3BioTEG/ | CENP-B pull-down |
| Cen_hairpin_R | AAAAGAAAGGTGTAACACACAAAGAAGTTTCT CAGAAAGCTTCTTTCTAGTTTTTATCTG | CENP-B pull-down |
